# Antibiotic resistance genes and mobile genetic elements removal from treated wastewater by sewage-sludge biochar and iron-oxide coated sand

**DOI:** 10.1101/2020.09.17.302018

**Authors:** David Calderón-Franco, Apoorva Seeram, Gertjan Medema, Mark C. M. van Loosdrecht, David G. Weissbrodt

## Abstract

Disinfection of treated wastewater in wastewater treatment plants (WWTPs) is used to minimize emission of coliforms, pathogens, and antibiotic resistant bacteria (ARB) in the environment. However, the fate of free-floating extracellular DNA (eDNA) that do carry antibiotic resistance genes (ARGs) and mobile genetic elements (MGEs) is overlooked. Water technologies are central to urban and industrial ecology for sanitation and resource recovery. Biochar produced by pyrolysis of sewage sludge and iron-oxide-coated sands recovered as by-product of drinking water treatment were tested as adsorbents to remove ARGs and MGEs from WWTP effluent. DNA adsorption properties and materials applicability were studied in batch and up-flow column systems at bench scale. Breakthrough curves were measured with ultrapure water and treated wastewater at initial DNA concentrations of 0.1-0.5 mg mL^-1^ and flow rates of 0.1-0.5 mL min^-1^. Batch tests with treated wastewater indicated that the adsorption profiles of biochar and iron-oxide coated sand followed a Freundlich isotherm, suggesting a multilayer adsorption of nucleic acids. Sewage-sludge biochar exhibited higher DNA adsorption capacity (1 mg g^-1^) and longer saturation breakthrough times (4 to 10 times) than iron-oxide coated sand (0.2 mg g^-1^). The removal of a set of representative ARGs and MGEs was measured by qPCR comparing the inlet and outlet of the plug-flow column fed with treated wastewater. ARGs and MGEs present as free-floating eDNA were adsorbed by sewage-sludge biochar at 85% and iron-oxide coated sand at 54%. From the environmental DNA consisting of the free-floating extracellular DNA plus the intracellular DNA of the cells present in the effluent water, 97% (sewage-sludge biochar) and 66% (iron-oxide coated sand) of the tested genes present were removed. Sewage-sludge biochar displayed interesting properties to minimize the spread of antimicrobial resistances to the aquatic environment while strengthening the role of WWTPs as resource recovery factories.

**Graphical abstract:** 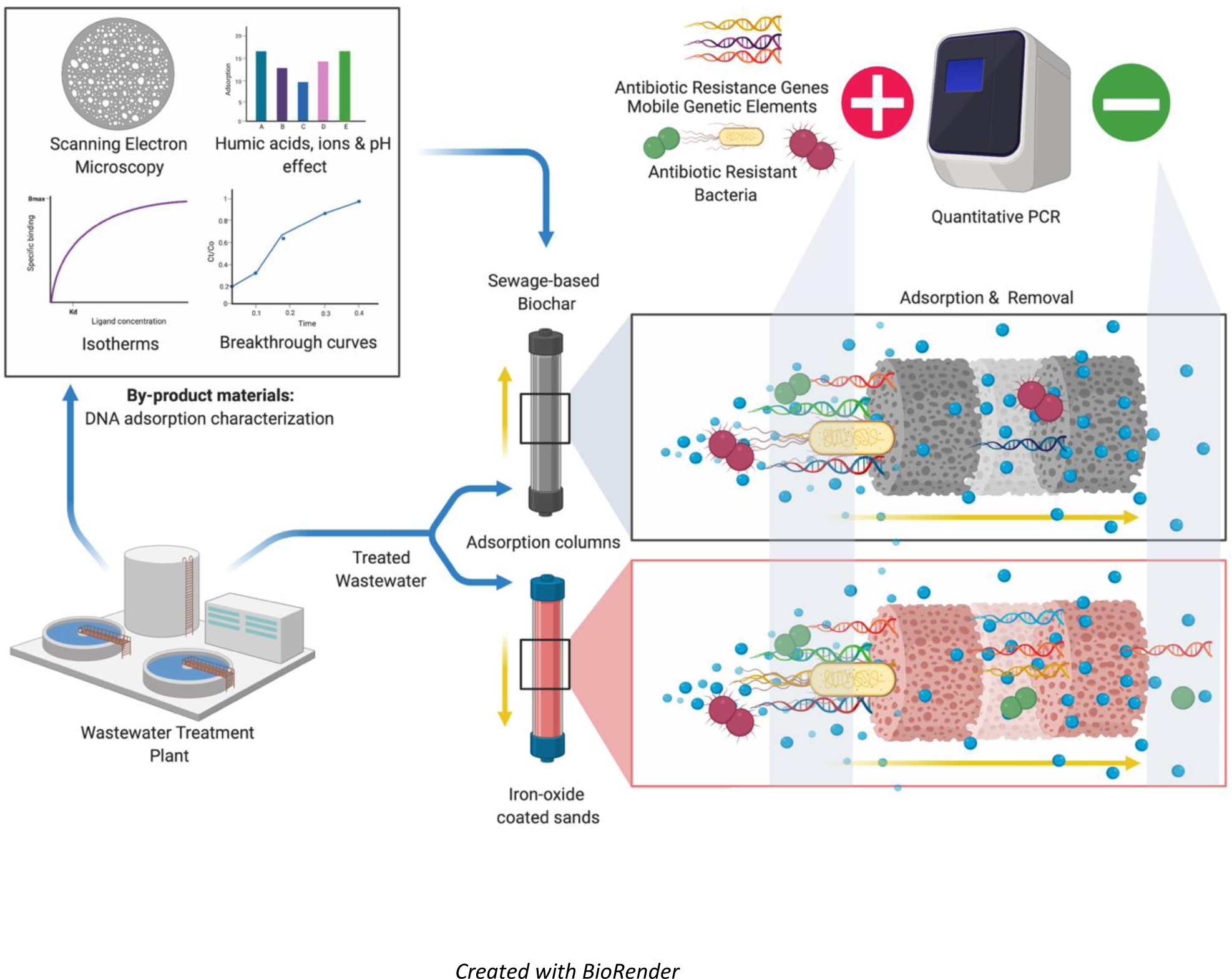

**Highlights:**

- Sewage-sludge biochar and iron oxide coated sands were tested to adsorb DNA and cells.
- Biochar removed 97% of genes tested from environmental DNA of unfiltered effluent.
- 85% of ARGs and MGEs of free-floating extracellular DNA were retained by biochar.
- Biochar is a WWTP by-product that can be re-used for public health sanitation.

## 1 Introduction

According to the UNICEF et al. (2019), about 785 million people in the world do not have access to potable water. An increase in population and living standards will create a scarcity on water resources availability (Shannon et al., 2008). New and alternate solutions to provide clean water are needed, such as wastewater reuse. The main problem with the reuse of wastewater is the final effluent quality. Different treatments are necessary based on the end use of the reclaimed water. Sustainable Development Goals (SDG 6.3) targets high water quality by reducing the use of hazardous materials and increasing the proportion of treated water, thus stimulating recycling and safe reuse.

Wastewater treatment plants (WWTPs) receive sewage from different sources such as households, public institutions such as hospitals or research centres, industries, urban runoff and agricultural streams (European Environment Agency, 2018). Conventional WWTPs are generally designed to remove pathogens, solids, organic matter and nutrients like nitrogen and phosphorus. They are not designed to treat a number of persistent xenobiotic compounds, which are often discharged with the effluent (McEachran et al., 2018). Only a few countries have adopted quality criteria in their water protection legislation to address the emissions and removal of micropollutants from wastewater, and their ecological impacts (Eggen et al., 2014; Weissbrodt et al., 2009). Both chemical and biological contaminants rise concerns.

Antimicrobial resistance (AMR) are important by the threat caused by the generation and spread of antibiotic-resistant pathogens across water bodies, soil and the food chain, impacting environmental, animal and human health (Pruden et al., 2006; Tripathi and Tripathi, 2017). The overuse and misuse of antibiotics affect the medical effectiveness in combating pathogenic organisms (C. Lee Ventola, 2019; Gould et al., 2015).

WWTPs play a central role in the recycling of water to the aquatic ecosystem. Located at the end of the sewer pipe, they face all pollutant loads emitted in the catchment area. New technical measures need to be developed to address micropollutants as well as antimicrobial resistance spreading. In contrast to chemicals, AMR can horizontally transfer, proliferate, and reproduce in biological environments. Because of the microbial diversity of the activated sludge, WWTPs are often hypothesized as hotspots for the horizontal gene transfer of AMR (Cacace et al., 2019)(Cacace et al., 2019)(Cacace et al., 2019; Rizzo et al., 2018).

Continuous release of antibiotics, ARB, and ARGs in wastewater may create a selective pressure leading to the development and proliferation of new ARG in activated sludge even if WWTPs prevents the sewer outflow to be a hotspot. However, the load is still high and there is a risk of human exposure and also horizontal gene transfer in receiving water bodies. ARB have been hypothesized to be mainly generated by horizontal gene transfer and take up of ARGs and mobile genetic elements (MGEs) by microorganisms (Karkman et al., 2018). Stress conditions and complex microbial communities may enhance the transformation of free-floating extracellular DNA (eDNA) fragments containing a variety of ARGs and MGEs (Lu et al., 2015; von Wintersdorff et al., 2016). MGEs have been shown to form the main component (65%) of eDNA free-floating in wastewater (Calderón-Franco et al., 2020).

When the resistome from 79 sites in 60 countries was examined, it was found that there are systematic differences in abundance and diversity of ARGs between continents and that ARGs abundance highly correlates with socioeconomic, health and environmental factors. Improving sanitation and health could potentially limit the global burden of antibiotic resistance (Hendriksen et al., 2019). A recent study of a set of six representative antibiotic resistant genes (ARGs) across more than 60 WWTPs of The Netherlands highlighted that the WWTPs do not amplify the release of these ARGs (Pallares-Vega et al., 2019). ARGs and MGEs, such as the class I Integron-integrase gene (*intI1*), have been reduced on average of a 1.76 log unit from the influents to the effluents. This however does not mean that no measure should be taken at WWTP level. ARGs persist in WWTP effluents as well as in river and lakes some km away from the effluent discharge (Czekalski et al., 2012). Tetracycline resistance genes like *tetO, tetQ, tetW, tetH, tetZ* have been quantified at an average of 2.5 x 10^2^, 1.6 x 10^2^, 4.4 x 10^2^, 1.6 x 10^1^ and 5.5 x 10^3^ gene copies mL^-1^ of chlorinated wastewater effluents, respectively (Al-Jassim et al., 2015).

Tertiary treatments like effluent disinfection using UV or chlorination do not suppress the release of ARGs in the environment, these genes can still be detected in disinfected effluents (Pazda et al., 2019). Disinfection can inactivate or select for ARB (Destiani and Templeton, 2019; Wu and Xu, 2019; Yuan et al., 2015), while the genes can remain and be released as extracellular DNA (eDNA) by cell lysis. Most studies on AMR did not consider the eDNA. The concentrations of free-floating eDNA measured from different wastewater samples (influent, activated sludge and effluent) ranged between 2.6 to 12.5 µg L^-1^ (Calderón-Franco et al., 2020a). This eDNA fraction can persist several months or even years in marine and soil matrices (Mao et al., 2014; Torti et al., 2015). It can attach to suspended particles, sand, clay and humic acids. However, sorption of eDNA on natural particles does not prevent its mobility and ability to transform into natural competent bacteria (Zhang et al., 2018).

ARGs that are not removed by neither physical, chemical nor biological treatment processes, being designated as persistent (Yang et al., 2014). Until now, different technologies can remove ARBs and ARGs from wastewater effluents. These can go from membrane bioreactor treatments (Kappell et al., 2018; Riquelme Breazeal et al., 2013) to coagulation (Li et al., 2017) or algal-based wastewater treatment systems (Cheng et al., 2020). The aforementioned methods however employ non-renewable materials or require consistent variations on the WWTP operation processes.

For the removal of free-floating extracellular DNA, the use of resources derived from the water cycle would be beneficial in the circular economy and resource recovery context (Van Der Hoek et al., 2016). Sewage sludge biochar produced by pyrolysis of activated sludge is an interesting recycling resource for soil amendment to immobilize heavy metals (Cd, Cu, Ni, Pb or As) and prevent environmental risk (Agrafioti et al., 2013). Iron-oxide coated sands are used to remove metals (As, Ni, Zn or Fe) from drinking water.

Here, in a sanitation and circular economy approach of WWTP, we assessed the reuse of by-products such as sewage-sludge biochar prepared by pyrolysis of dewatered sewage sludge and iron-oxide coated sands reclaimed from drinking water processing for their adsorption capacity of environmental and extracellular DNA including ARGs and MGEs as well as ARB from secondary wastewater effluents. First, the isotherms and mechanisms governing DNA adsorption onto these by-product materials were studied both in batch and up-flow column experiments under both synthetic aqueous conditions and real effluent wastewater. Second, the fate and abatement of the environmental DNA (including both the intracellular and extracellular DNAs) and of the free-floating eDNA from samples of real treated wastewater effluent were analyzed across fixed beds of sewage-sludge biochar and iron-oxide coated sands. We show that sewage-sludge biochar can efficiently remove both the microorganisms and the free-floating genes.

## 2 Material and Methods

### 2.1 Sampling from the effluent of a wastewater treatment plant

Biological samples of effluent water were collected from the urban wastewater treatment plant (WWTP) Harnaschpolder (Waterboard Delfland, The Netherlands) operated for full biological nutrient removal. Effluent water was collected at the outlet of the tertiary treatment of WWTP Harnaschpolder. Three biological replicates were collected in three different days. A total of 1000 mL of treated water per replicate was collected. All samples were processed in a timeframe of less than 4 h.

### 2.2 Dewatered sewage-sludge and iron-oxide coated sand collection

Dewatered sewage sludge to produce biochar was also collected from Harnaschpolder (Waterboard Delfland, The Netherlands). The dewatered sewage sludge samples were brought to the lab within 1 hour and stored at 4° C until further analysis.

An amount of 2 kg of iron oxide coated sand (1-4 mm) was reclaimed from the sand-filtration unit at AquaMinerals® (https://aquaminerals.com/home/), a water sanitation company giving a second life to the resources from WWTP. Iron-oxide coated sands received were crushed and sieved under 600 µm and stored at room temperature for further experiments.

### 2.3 Sewage-sludge biochar production by pyrolysis

The sewage-sludge before pyrolysis was characterized by moisture content and volatile matter. The moisture content was performed at 11 ± 1°C and volatile matter at 500 ± 50°C until constant weight was achieved. Sewage sludge biochar was produced following the procedure as described in Agrafioti *et al*. (2013). Briefly, dewatered sludge was heated in a muffle furnace at 600°C under a controlled flow of nitrogen. The samples were kept for 30 minutes residence time. The cooled samples were then crushed and sieved at 150 µm pore size. The methodology for determining the properties of raw material used for the production of biochar is followed by the study done by Agrafioti *et al*. (2013). Sewage-sludge biochar produced was stored at room temperature for further experiments. The sewage-sludge biochar yield was determined as the ratio of the produced dry mass of sewage-sludge biochar (after pyrolysis) to the dry mass of sewage-sludge (before pyrolysis).

### 2.4 Adsorbents chemical characterization

The inorganic chemical composition of sewage-sludge biochar was characterized using Panalytical Axios Max WD-XRF spectrophotometer and the data was evaluated with SuperQ5.0i/Omnian software. Carbon (C), oxygen (O) and nitrogen (N) could not be measured by WD-XRF spectrophotometer. For performing such analysis, the elemental composition of carbon (C), oxygen (O), hydrogen (H), nitrogen (N) and sulphur (S) from sewage sludge biochar and iron (Fe) and silica (Si) from iron-oxide coated sand were analyzed by an elemental analyser (Mikrolab Kolbe, Germany).

### 2.5 Adsorbent surface area and pore size determination

Surface area and pore size of the adsorbents were calculated using a nitrogen gas adsorption analyzer (Micrometrics Gemini VII 2390 Surface area analyser, USA). Before analysis, 5 g of the by-product materials were previously degassed under vacuum at 150°C (under N2 flow) during 1 h to eliminate moisture and gasses. Then, 0.5 g of the by-product materials were subsequently introduced in the test tube with liquid nitrogen. Finally, N2 isotherms were measured at -196°C. Surface area was calculated according to Brunauer - Emmett - Teller (BET) method (Naderi, 2015), which incorporates a multilayer coverage. Barret, Joyner and Halenda (BJH) method was used to determine the pore size using Kelvin equation of pore filling, where a cylindrical pore geometry was assumed (Villarroel-Rocha et al., 2014).

### 2.6 Iron-oxide coated sands valence state determination

Valence state of iron (Fe) in iron-oxide coated sand was determined by Mössbauer Spectroscopy in order to know the oxidation state. A conventional constant – acceleration spectrometer the absorption spectra for 300 K and at 4.2 K with a sinusoidal velocity spectrometer, using a ^57^Co (Rh) source were measured. A velocity calibration curve using an α-Fe foil at room temperature was done.

### 2.7 Scanning electron microscopy

The surface observation of sewage-sludge biochar and iron-oxide coated sand was analyzed by scanning electron microscopy (SEM), using JOEL model JSM-6010LA. For the observation of the above samples, we used the same SEM magnifications (5K, 10K, 15 means a thousand), same acceleration voltage (5-15 KeV).

### 2.8 DNA template and chemicals

UltraPure™ Salmon sperm DNA solution (Thermo Fisher Scientific, USA) for batch and column studies was used as a representative model for eDNA. Salmon sperm DNA is double stranded, sheared to an average size of ≤2,000 bp. The purity of DNA was assessed by UV light absorbance at 260 and 280 nm (A260/280 > 1.8). The chemical and other reagents were obtained from Sigma Aldrich (USA). In order to assess the effect of different ions and humic acids on DNA adsorption, stock solutions of 1 M of salts (NaCl, CaCl2.2H2O, MgCl2.6H2O) and 800 mg L^-1^ humic acids (CAS No. 1415-93-6) were prepared by adding in ultrapure water (Sigma Aldrich, UK).

### 2.9 Determination of the adsorption equilibria of the adsorbents

To find the equilibrium time required for the adsorption by sewage-based biochar and iron-oxide coated sand, batch experiments were done in 6 apothecary glass bottles of 100 mL, with a working solution of 5 mL. Initial salmon sperm DNA concentration of 100 µg mL^-1^ was added. The mass of the adsorbent (sewage-sludge biochar or iron-oxide coated sand) was added increasing from 0 to 100 mg mL^-1^ in separate bottles and in triplicates. The bottles were kept for mixing at room temperature continuously for 24 hours. 0.5 mL of the sample was taken every 1 h and centrifuged at 13000 x G for 20 min.

### 2.10 Effect of pH, ionic strength and humic acids content on DNA adsorption

To evaluate the influence of pH on adsorption, experiments were done in 10 mM Tris-HCl buffer with initial DNA concentration of 20 and 100 µg mL^-1^ at pH 5, 7 and 9. The cation species effect on adsorption was also studied at initial DNA concentration of 100 µg mL^-1^ in the presence of 0 – 60 mM Na^+^ (as NaCl), Mg^2+^ (as MgCl2), and Ca^2+^ (as CaCl2) at pH 7, respectively. Competition with organic matter on DNA adsorption was investigated using humic acid to represent natural organic matter. The experiments were done in the presence of 0-100 mg L^-1^ humic acids at pH 7. In order to maintain constant pH, the effect of cation species and competition with humic acids was done in Tris-HCl buffer.

### 2.11 Quantification of adsorbed DNA

The DNA concentration in the supernatant was measured by UV spectrometry at a wavelength of 260 nm (BioTek, Gen5 plate reader, USA). 96-well UV flat-bottom plates (Greiner UV Star 96, Germany) were used for measuring the absorbance. The DNA concentration after incubation with humic acids was measured with Qubit® dsDNA assays (Thermo Fisher Scientific, USA) (Leite et al., 2014). The amount of DNA adsorbed onto the adsorbent at equilibrium was calculated using **equation 1**.

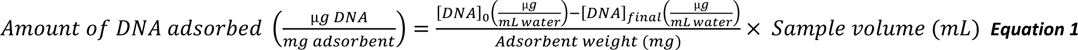

### 2.12 Adsorption isotherms and sorption efficiency

Two adsorption isotherm models were used in these experiments: Langmuir and Freundlich isotherms. The Langmuir describes adsorption of adsorbate (DNA in this case) molecules by assuming that it behaves as an ideal gas at isothermal conditions. Adsorption is supposed to happen onto homogeneous solid surfaces that exhibit one adsorption site. The Freundlich isotherm is an empirical relation between the concentration of a solute on the surface of an adsorbent (by-products) to the concentration of the solute in the liquid with which it is in contact. The Langmuir model assumes that at maximum coverage, there is only a monomolecular layer on the surface. It means that there is no stacking of adsorbed molecules. The Freundlich isotherm does not have this restriction.

Both Freundlich and Langmuir isotherms were used to describe DNA adsorption from the solution onto the adsorbent. The Freundlich isotherm is expressed by **equation 2.**

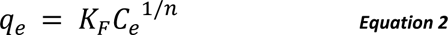

where qe (mg g^-1^) is the amount of solute adsorbed, Ce (mg L^-1^) is the equilibrium adsorbate concentration, KF (mg g^-1^) is the Freundlich constant related to the adsorption capacity and n (without units) is the heterogeneity factor and adsorption favourability.

The Langmuir isotherm is expressed as **equation 3.**

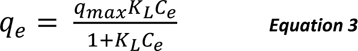

where qe (mg g^-1^) is the amount of adsorbate adsorbed, Ce (mg L^-1^) is the equilibrium adsorbate concentration, qmax (mg g^-1^) is the maximum monolayer adsorption capacity and KL (L mg^-1^) is the Langmuir empirical constant related to the heat of adsorption. KL represents the adsorption affinity of the adsorbate onto the adsorbent

### 2.13 Fixed-bed column characteristics and operation

Continuous-flow adsorption experiments were conducted in chromatography glass columns (1 cm inner diameter and 15 cm column). At the bottom inside of the column, 0.5 g of glass beads (250 – 300 µm) were placed for homogeneous flow through the column distribution. The columns were packed with known quantities of adsorbents: 3 g for sewage-sludge biochar and 4 g for iron-oxide coated sand. The column was fed with an aqueous solution of salmon sperm DNA of known concentration using an HPLC liquid chromatography pump (Shimadzu LC-8A, USA) employing an upward flow. Samples (200 µL) were collected at the outlet of the column in intervals of 15 min. DNA concentration was measured spectrophotometrically until the ratio of Ct/Co reached a constant value, where Co is the column inlet stable DNA concentration and Ct is the column outlet DNA concentration. When the Ct/Co ratio was close to 1, it meant that the column was saturated of DNA.

To determine the adsorption breakthrough curves, the inlet DNA concentrations were modified from 0.1 mg mL^-1^, 0.3 mg mL^-1^ and to 0.5 mg L^-1^ at a fixed flow rate of 0.1 mL min^-1^. The flow rates were modified from 0.1 mL min^-1^, 0.3 mL min^-1^ and 0.5 mL min^-1^ at a fixed inlet concentration of 0.3 mg L^-1^.

### 2.14 Column data analysis

Performance of a fixed bed column can be explained by the breakthrough curves. The amount of time needed for breakthrough and the shape of the curve gives the dynamic behavior of the column. Breakthrough curves are expressed as Ct/Co versus time (Han et al., 2009; Rouf and Nagapadma, 2015). The breakthrough point is usually defined as the point when the ratio between influent concentration, C0 (mg L^-1^) and outlet concentration, Ct (mg L^-1^) becomes 0.05 to 0.90 (Chowdhury et al., 2015). The total capacity of the column (qtotal in mg) gives the maximum amount of DNA that can be adsorbed and is calculated by the area under the breakthrough curve given by **equation 4** (Chen et al., 2012; Han et al., 2009; Rouf and Nagapadma, 2015)

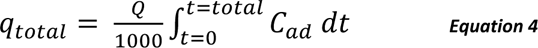

where Q is the flow rate (mL min^-1^); t=total is the total flow time (min); Cad is the adsorbed DNA concentration (Co – Ct) (mg L^-1^).

The equilibrium DNA uptake or maximum adsorption capacity of the column qeq(exp) (mg g^-1^) is calculated by **equation 5**:

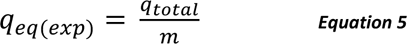

where m is the dry weight of the adsorbent in the column (g).

### 2.15 Modelling of the fixed-bed column

The data generated from the study were fitted with Thomas’ and Yoon-Nelson’s models for column modelling as in Chatterjee et al. (2018).

In order to design an adsorption column, prediction of breakthrough curve and adsorbent capacity for the adsorbate under certain conditions is required. Data obtained from the experiments can be used for designing a prospective full-scale column operation. In this research, data from column studies have been analyzed using the Thomas’ model.

Thomas’ model was used to estimate the absorptive capacity of the adsorbent. The expression for the Thomas model is given in **equation 6**.

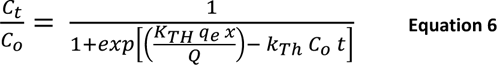

where *k*_TH_ (mL min ^-1^ mg^-1^) is the Thomas model constant; *q*_*e*_(mg g^-1^) is the predicted adsorption capacity; *x* is the mass of adsorbent (g); *Q* is the flow rate (mL min^-1^); *C*_*o*_is initial DNA concentration (mg L^-1^); *C*_*t*_ is the effluent concentration at time t (mg L^-1^).

Yoon-Nelson’s model was used to predict the time of run before regeneration or replacement of the column becomes necessary. It is a very simple model to represent the breakthrough curve as it does not require any data about the characteristics of the system and the physical properties of the adsorbent (Rouf and Nagapadma, 2015). Yoon-Nelson’s model can be expressed as **equation 7**.

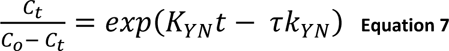

where *K*_*YN*_ (min^-1^) is the rate constant; τ (min) is the time required for 50% adsorbate breakthrough.

### 2.16 Hydraulic residence time in the column

Hydraulic residence time distributions in the columns filled with sewage-sludge biochar and iron-oxide coated sand was performed by using NaCl salt tracer. The columns were filled with sewage-sludge biochar or iron-oxide coated sand with a bed height of 10 cm and a flow rate of 1 mL min^-1^ was used. A concentration of 60 mM NaCl salt solution was pulse dosed into the column by using an HPLC liquid chromatography pump (Shimadzu LC-8A, USA). The concentration changes in the effluent was measured by electrical conductivity using PRIMO 5 Microprocessor Conductivity Meter (Hanna instruments, USA), as a function of time by taking samples of 0.5 mL.

### 2.17 DNA extraction before and after loading adsorption columns

Two types of DNA were extracted: (*i*) free-floating eDNA from filtered treated wastewater and (*ii*) environmental DNA (intracellular and extracellular DNA) from unfiltered treated wastewater before loading the column and in the column eluents. The isolation of free-floating extracellular DNA and of intracellular DNA has been described by Calderón-Franco et al. (2020).

For extracting the free-floating extracellular DNA, 1000 mL of raw effluent samples were filtered sequentially through 0.44 µm and 0.22 µm polyvinylidene fluoride (PVDF) (Pall Corporation, USA) membranes and further processed for isolating free-floating extracellular DNA. 1000 mL of filtered effluent sample (containing the free-floating extracellular DNA) was loaded in a positively charged 1 mL diethylaminoethyl cellulose (DEAE) column (BIA Separations, Slovenia) at a speed of 0.6 mL min^-1^ after equilibration in order to keep the pressure below 1.8 MPa (pressure limit for the 1 mL column).

Buffers and solutions used for equilibrating, eluting, regenerating, cleaning and storing the column are the following: Equilibration buffer consisted on 50 mM Tris, 10 mM EDTA at pH 7.2. Elution buffer consisted on 50 mM Tris, 10 mM EDTA, 1.5 M NaCl at pH 7.2. Regeneration buffer consisted on 50 mM Tris, 10 mM EDTA, 2 M NaCl, pH 7.2. Cleaning solution consisted on 1 M NaOH and 2 M NaCl. Storage solution consisted of 20% Ethanol.

Column preparation and processing were done according to manufacturer’s instructions. Elution was done at a speed of 1 mL min^-1^ with elution buffer and eluent was tracked over time with an HPLC photodiode array detector (Waters Corporation, USA) recording the UV-VIS absorption at wavelength 260 nm. The eluent was further precipitated with ethanol (Moore and Dowhan, 2002) to obtain the raw free-floating extracellular DNA sample. Precipitated raw free-floating extracellular DNA was incubated with 0.85 g L^-1^ proteinase K (Sigma-Aldrich, UK) during 2 hours and enzymatic reaction was stopped at 50°C for 10 minutes. Protein-digested raw extracellular DNA sample was finally purified using GeneJET NGS Cleanup Kit (Thermo Scientific, USA). Obtained free-floating extracellular DNA samples were stored at -20°C for further analysis.

Environmental DNA (extracellular and intracellular DNA) was extracted from the unfiltered water sample before loading the column and the effluent using the DNeasy kit Power Water (Qiagen, The Netherlands) as per the instructions given by the manufacturer. The experiments were performed in triplicates. The extracted DNA from filtered and unfiltered samples were quantified by fluorometry using Qubit® (Thermo Fisher Scientific, USA).

### 2.18 Quantification of ARGs and MGE by quantitative polymerase chain reaction

The 16S rRNA gene was selected as a proxy to quantify total bacteria. The genes analyzed by qPCR were chosen and modified from a selection panel of antibiotic resistance genes already used for wastewater samples (Pallares-Vega et al., 2019). Standards for qPCR were generated from ResFinder (https://cge.cbs.dtu.dk/services/ResFinder/), a curated antimicrobial resistance genes database. The chosen ARGs confer resistance to antibiotics with the highest consumption in The Netherlands: macrolides (*ermB*), sulfonamides (*sul1* and *sul2*), fluoroquinolones (*qnrS*) and extended-spectrum β-lactamase (*blaCTXM*) **(Table 1).** Moreover, a gene assessing the presence of MGE was included: *intI1,* an integrase of class I Integron, known to be jumping genes responsible of driving horizontal gene transfer phenomena (Ma et al., 2017).

**Table 1.** Details of the panel of universal 16S rRNA gene, antibiotic resistant genes (ARGs), and mobile genetic element (MGE) used for the study. The selection was based on the work of Pallarés-Vega et al. (2019) as representative genes for The Netherlands.

| Group | Gene | Function |
| --- | --- | --- |
| All Bacteria | 16S ribosomal RNA gene | Normalization to the concentration of bacteria |
| ARGs | *ermB* (Erythromycin resistance) | Resistance to macrolides |
|  | *sul1* (Sulfonamide resistant dihydropteroate synthase 1) | Resistance to sulfonamides |
|  | *sul2* (Sulfonamide resistant dihydropteroate synthase 2) | Resistance to sulphonamides |
|  | *qnrS* (Fluoroquinolone resistance) | Resistance to quinolones |
| | *bla<sub>CTXM</sub>* (Cefotaxime-hydrolyzing β-lactamase resistance) | Resistance to extended spectrum β-lactams |
| MGE | *int11* (Class 1 Integrase) | Integrase of type 1 integrons |

The selected genes for the analysis are shown in **Table 1**. Standards, primers and reaction conditions used in this study are listed in the supplementary material and **tables S1 and S2**.

## 3 Results and Discussion

### 3.1 By-products materials show a porous morphology optimal for adsorption experiments

Two by-product materials were selected to investigate the removal of DNA fragments by adsorption. Sewage-sludge biochar was produced at high temperature (600°C) from excess sludge collected from a WWTP. Reclaimed iron-oxide coated sand was obtained from a drinking water treatment plant. **Figure 2** shows scanning electron microscopy micrographs of sewage-sludge biochar **(Fig. 2a-b)** and iron-oxide coated sand **(Fig. 2c-d)**. The surface morphologies of both by-products were highly heterogeneous and structurally complex, with many pores of different diameters. No fibers nor other debris structures were observed in any of the materials.

**Figure 1.**
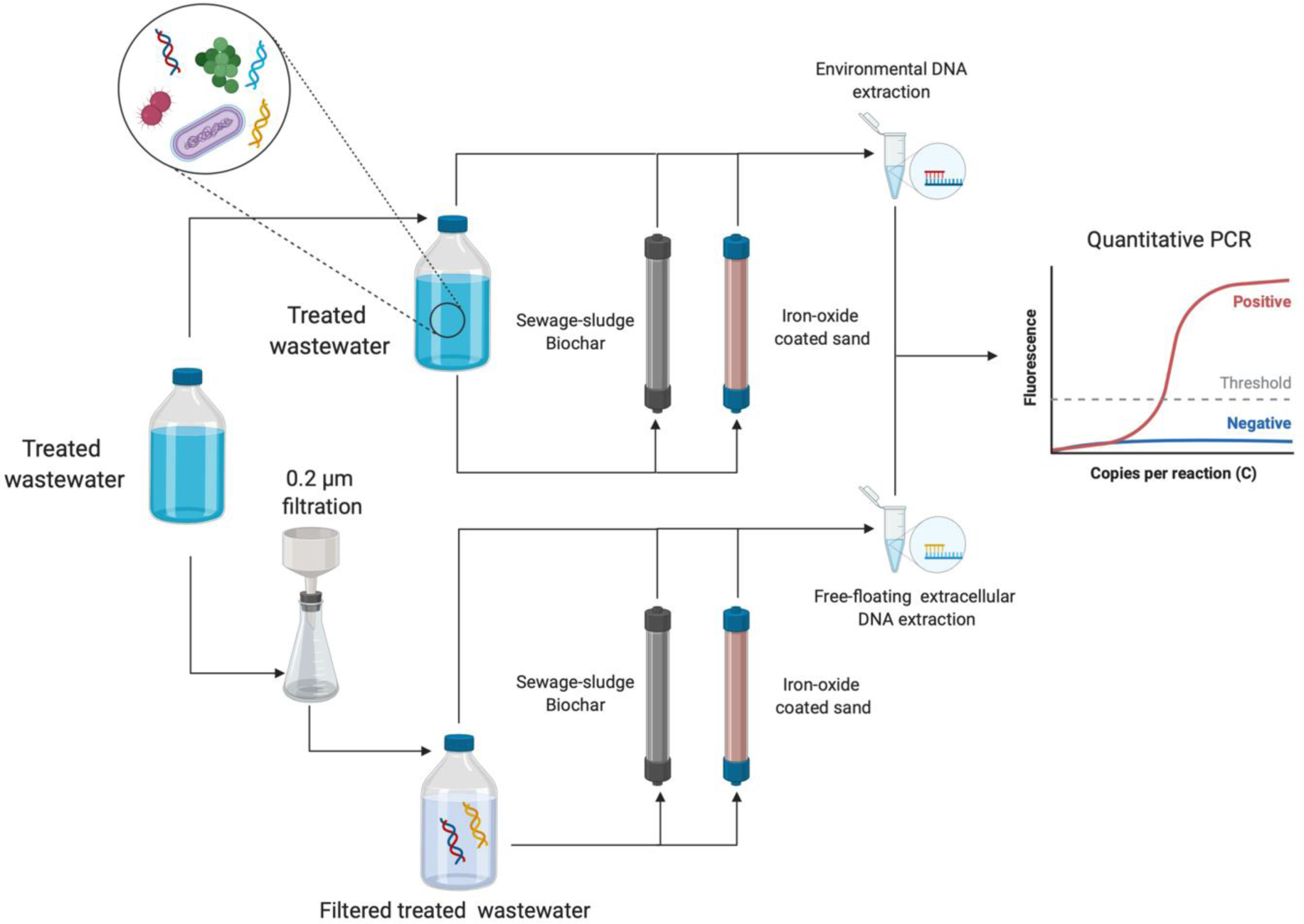
Schematic representation of the experimental adsorption column setup used and the quantitative PCR as analytical procedure applied. The aim was to study the adsorption and removal of antibiotic resistance bacteria, antibiotic resistance genes and mobile genetic elements by using sewage-sludge biochar and iron oxide-coated sand from treated effluent wastewater. Detail of material and setup used can be found in **Figure** S2. Created with *BioRender*.

**Figure 2.**
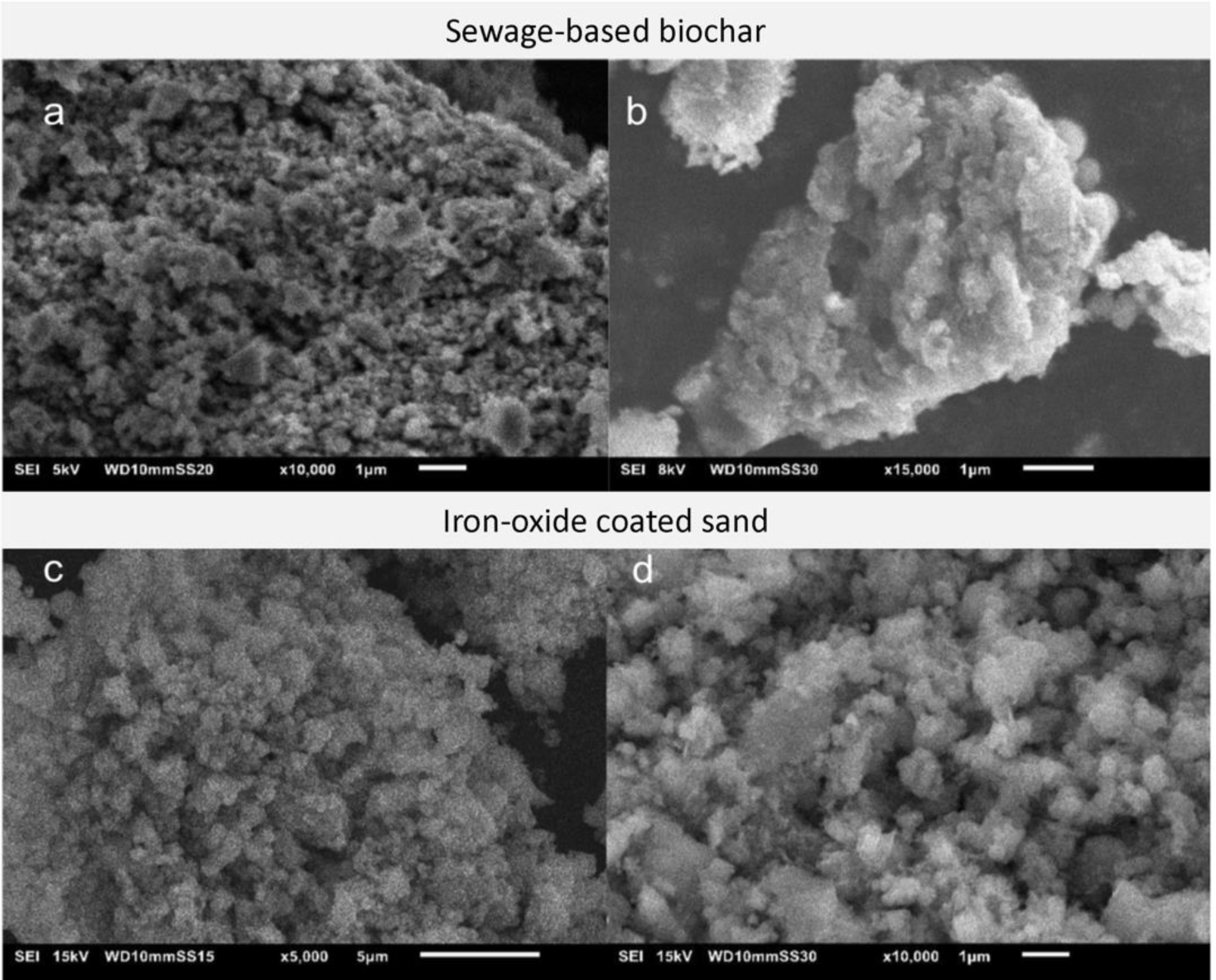
Scanning electron microscopy images of sewage sludge biochar at magnifications of **(a)** 10.000x and **(b)** 15.000 x and iron oxide coated sand at **(c)** 5.000x and **(d)** 10.000x.

The results of the Brunauer-Emmett-Teller (BET) specific surface area analyses are provided in **table 2.** Sewage-sludge biochar surface area was lower than for iron-oxide coated sand: 32.4 vs 164.9 m^2^g^-1^. Ash filling, which blocks access to the biochar micropores, could explain why specific surface areas in biochar are lower than in other materials (Song and Guo, 2012). Méndez *et al*. (2013) got similar BET surface area values and pore diameter with biochar pyrolyzed at 600°C from sewage sludge: 37.18 m^2^ g^-1^ and 9.46 nm, respectively.

**Table 2.** Physicochemical properties and the elemental composition of sewage-sludge biochar pyrolyzed at 600°C and iron oxide coated sand.

| Material | Sewage-based biochar | Iron-oxide coated sand |
| --- | --- | --- |
| Yield (%) | 38.6 | - |
| Water content (%) | 78.9 | - |
| Volatile matter (%) | 68.1 | - |
| Surface area (m<sup>2</sup>g<sup>-1</sup>) | 32.4 | 164.9 |
| Pore diameter (nm) | 10.1 | 6.2 |
| pH | 8.0 | 8.2 |
| Elemental composition analysis |  |  |
| C (%) | 14.1 | 2.07 |
| H (%) | 1.7 | 3.28 |
| O (%) | 37.4 | 43.7 <sup>a</sup> |
| N (%) | 1.9 | 0.01 |
| S (%) | 19.5 | - |
| P (%) | 7.7 | - |
| Ca (%) | 4.3 | - |
| Mg (%) | 5.6 | - |
| Fe (%) | 3.9 | 27.3 |
| Si (%) | 3.2 | 23.61 |
| Al (%) | 1.3 | - |

<sup>a</sup> Oxygen content estimation.

The iron-oxide coated sand had a specific surface area of 164.9 m^2^ g^-1^. Sharma (2001) described the iron-oxide coated sand specific surface area from 12 different Dutch drinking water plants in the range of 5.4 to 201 m^2^ g^-1^. They suggested that the BET surface area of iron-oxide coated sand could mainly depend on their residence time (from months to years) in the drinking water treatment plant.

The sewage-sludge biochar production yield of 0.39 g (dried sewage-sludge biochar) g^-1^ (dried sewage-sludge) matches other studies (Roberts et al., 2017; Tarelho et al., 2019). The biochar production yield has an important impact on its industrial applicability. It highly varies from the pyrolysis temperature used (Daful and R Chandraratne, 2018). The low sewage-sludge biochar production yields here obtained could be caused by the high pyrolysis temperatures that may release higher amounts of volatile matter (Titiladunayo et al., 2012).

The chemical properties of the materials are presented in **Table 2**. The sewage-sludge biochar was mainly composed of oxygen, sulphur and carbon with traces of other elements like phosphorus, magnesium or calcium bioaccumulated during wastewater treatment. Iron-oxide coated sands mainly comprised iron, oxygen and silica. The valence state of iron in the iron oxide coated sand was determined by Mössbauer spectroscopy, resulting in mainly Ferrihydrite (III) particles **(Fig. S1; Table S3).**

**Table 3.** Freundlich and Langmuir parameters on DNA adsorption with both sewage-sludge biochar and iron-oxide coated sand materials in treated wastewater matrix.

| Adsorbent | Type of water | Freundlich | | | Langmuir | | Correlation coefficient $R^2$ | |
| --- | --- | --- | --- | --- | --- | --- | --- | --- |
| | | $K_F$<br>($\text{mg g}^{-1}$) | $1/n$ | $n$ | $q_{\text{max}}$<br>($\text{mg g}^{-1}$) | $K_L$<br>($\text{L mg}^{-1}$) | Freundlich | Langmuir |
| Sewage-sludge biochar | Treated wastewater | 1.0 | 0.19 | 5.16 | 2 | 0.61 | 0.99 | 0.92 |
| Iron-oxide coated sand | Treated wastewater | 0.21 | 0.61 | 1.63 | 5.8 | 0.01 | 0.93 | 0.89 |

### 3.2 Adsorption of nucleic acids by sewage-sludge biochar and iron-oxide coated sand seems to follow a multilayer Freundlich isotherm

Salmon sperm DNA was used as a synthetic DNA template to analyze the adsorption equilibrium of the by-products. The adsorption equilibrium time was reached after 2 h for sewage-sludge biochar **(Figure S3a)** and after 5 h for iron-oxide coated sand **(Figure S3b)**. Sewage sludge biochar showed higher adsorption capacities than iron-oxide coated sand. Both Langmuir and Freundlich isotherms were used to fit the data. **Figure 3a** and **3b** showed the different adsorption isotherm curves of salmon sperm DNA on sewage-sludge biochar and iron-oxide coated sand, respectively. **Table 3** summarizes the different estimated adsorption parameters for both materials in treated wastewater. For the Langmuir equation, the maximum DNA adsorption capacity (qmax) by sewage sludge biochar was 2.0 mg g^-1^ and iron oxide coated sand was 5.8 mg g^-1^. The Langmuir KL (L mg^-1^) constant was higher in sewage-sludge biochar (0.61 L mg^-1^) than in iron-oxide coated sand (0.01 L mg^-1^. The Freundlich equation showed a different trend of adsorption of DNA onto the tested adsorbents. Sewage-sludge biochar had a higher Freundlich KF (1 mg g^-1^) constant than iron oxide coated sand (0.21 mg g^-1^). This result was contrary to what could be observed in the maximum adsorption capacity when Langmuir parameters were analyzed.

**Figure 3.**
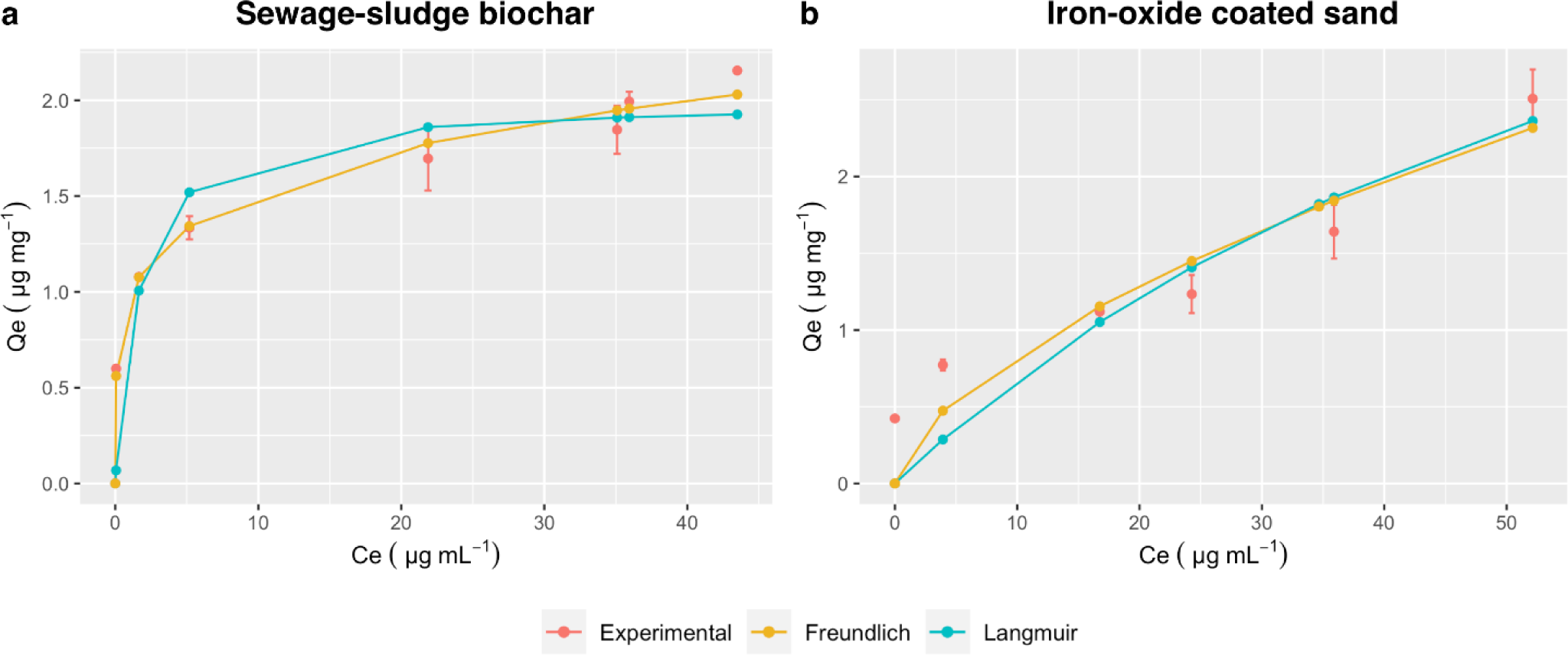
Freundlich and Langmuir isotherms models fitted from experimental data on **(a)** sewage sludge biochar and **(b)** iron oxide coated sand. In graph: Equilibrium salmon sperm DNA concentration (Ce) and the amount of salmon sperm DNA adsorbed per unit of adsorbent (Qe).

Comparing the correlation coefficient, the adsorption profiles can be better described by the Freundlich isotherm model (r^2^ = 0.99 and 0.93 for sewage-sludge biochar and iron-oxide coated sand, respectively) suggesting a multilayer adsorption with heterogeneous distribution of active adsorption sites rather than a Langmuir monolayer isotherm (r^2^ = 0.92 and 0.89 for sewage-sludge biochar and iron-oxide coated sand, respectively). For the adsorption to be thermodynamically favorable, the constant for a given adsorbate and adsorbent at a particular temperature “n” should lie between 1 and 10 (Desta, 2013). Values of “n” obtained from both of our tested adsorbents lied within the range **(Table 3)**. In terms of adsorption capacity, sewage-sludge biochar was found to be a more effective adsorbent for remediating potential xenogenetic elements.

Overall, biochar showed higher affinity towards nucleic acids. However, it is important to highlight that the binding mechanisms might be quite different between the materials tested. Likely, iron binds to the phosphate groups from DNA whereas on biochar, hydrophobic forces may be driving the interaction.

### 3.3 Effects of pH and coexisting anions

The influence of pH on DNA adsorption by sewage-sludge biochar and iron-oxide coated sand was analysed in **Figure 4.** The initial concentrations of DNA were 20 mg L^-1^ **(Figure 4a)** and 100 mg L^-1^ **(Figure 4b).** There was no effect observed of pH on the adsorption capacity of salmon sperm DNA for the tested materials.

**Figure 4.**
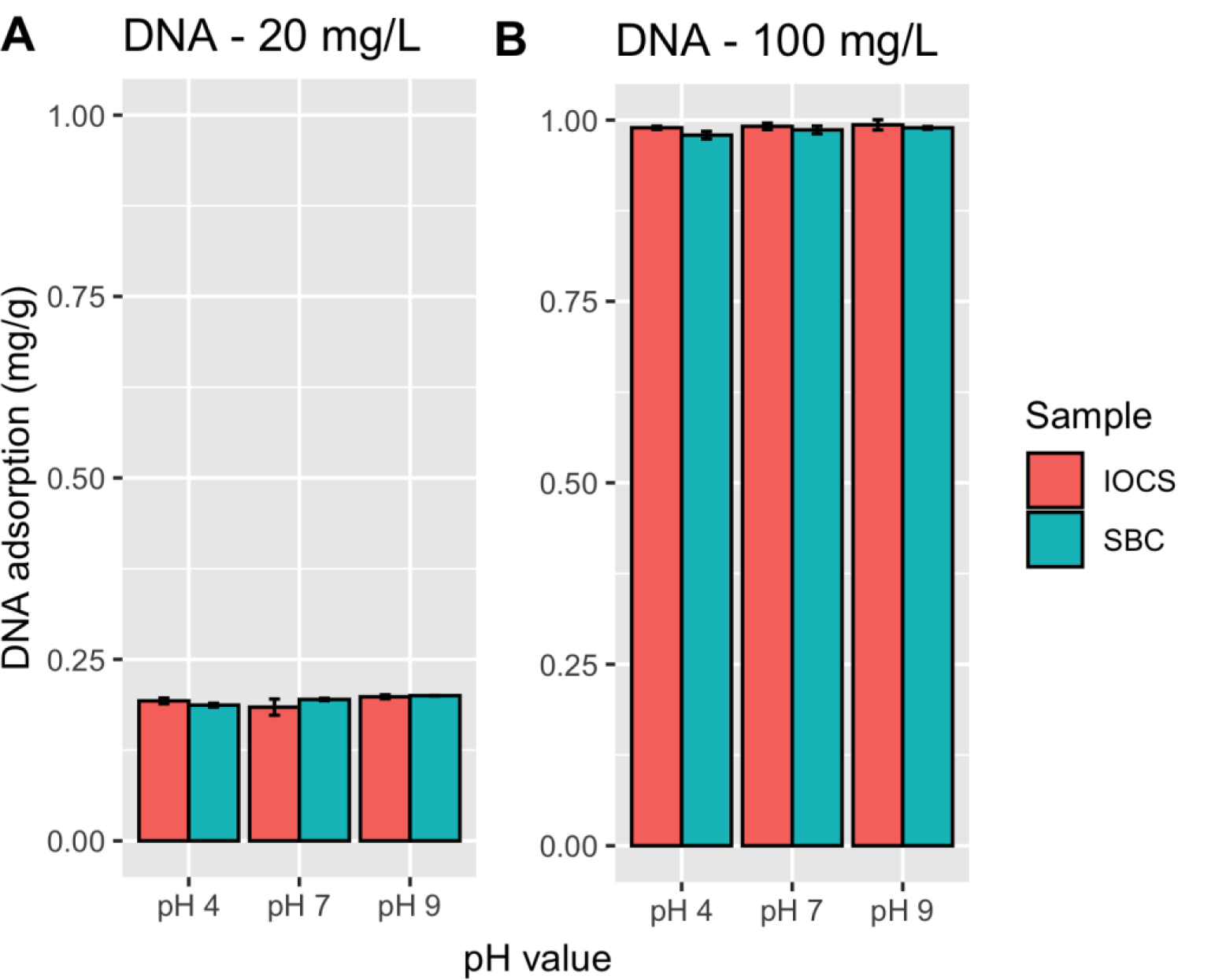
The influence of pH on DNA adsorption (mg DNA g material ^-1^) by sewage-sludge biochar (SBC) and iron-oxide coated sands (IOCS). The initial concentration of DNA was 20 mg L^-1^ (A) and 100 mg L^-1^ (B), respectively. No significant difference (p<0.05) among treatments were observed. DNA adsorption stands for mg of DNA per g of adsorbent added.

It has been described that organic clays, montmorillonite and biochar adsorb more DNA under acidic (pH < 5) than alkaline (pH > 9) conditions (Cai et al., 2006c; Saeki and Kunito, 2010). The biochar surface displays an increase of its negative charge when pH values range from 3 to 7 (Yuan et al., 2011). The isoelectric point of DNA is about pH 5 (Cai et al., 2006b). DNA molecules are negatively charged due to the phosphate groups when pH is above the isoelectric point. The phosphate groups from DNA at low pH < 2 are mostly present as a neutral species. In order to avoid electrostatic repulsion, sewage-sludge biochar and DNA should not be negatively charged simultaneously. This means that at pH lower than 4, both biochar surfaces and DNA phosphate groups are not negatively charged, decreasing repulsion between them. Thus, it should increase the adsorption capacity. However, in our case theory does not align with the experiments. Wang *et al*. (2014) have already observed that pH shifts on willow wood biochar did not display any significant influence on nucleic acids adsorption.

Adsorption of ortho-phosphate onto iron-oxide coated sand increase as the pH decreases, following an anionic adsorption behavior (Huang et al., 2014). In our case, pH shifts did not show any significant effect on DNA adsorption by the phosphate groups.

**Figure 5** presents the influence of ionic conditions on the adsorption of DNA to sewage-sludge biochar and iron-oxide coated sand at pH 7, at an initial DNA concentration of 100 mg L^-1^. The addition of the different ions (Ca^2+^, Mg^2+^ and Na^+^) in the concentrations range of 1 – 60 mmol L^-1^ had no significant effect (p>0.05) on the adsorption of salmon sperm DNA onto sewage-sludge biochar and iron-oxide coated sand (supplementary **table S4).** The only exception was the addition of 60 mmol Mg^2+^ L^-1^ on the sewage-sludge biochar material that did show a significant increase (0.89 mg DNA g^-1^, 32.6%) on the adsorption capacity (p<0.05). Ion bridges and charge neutralization are the supposed main mechanisms increasing DNA adsorption by cations addition (Cai et al., 2006a; Nguyen and Chen, 2007). However, DNA adsorption onto these materials did not seem to be driven by electrostatic interactions. For this reason, it is hypothesized that hydrophobic interactions or iron-phosphate interactions can play a role in the DNA adsorption onto the tested materials. It has been described that higher pyrolysis temperatures (> 600 °C) significantly increase DNA adsorption efficiency onto biochar due to higher surface area and hydrophobicity (Dai et al., 2017).

**Figure 5.**
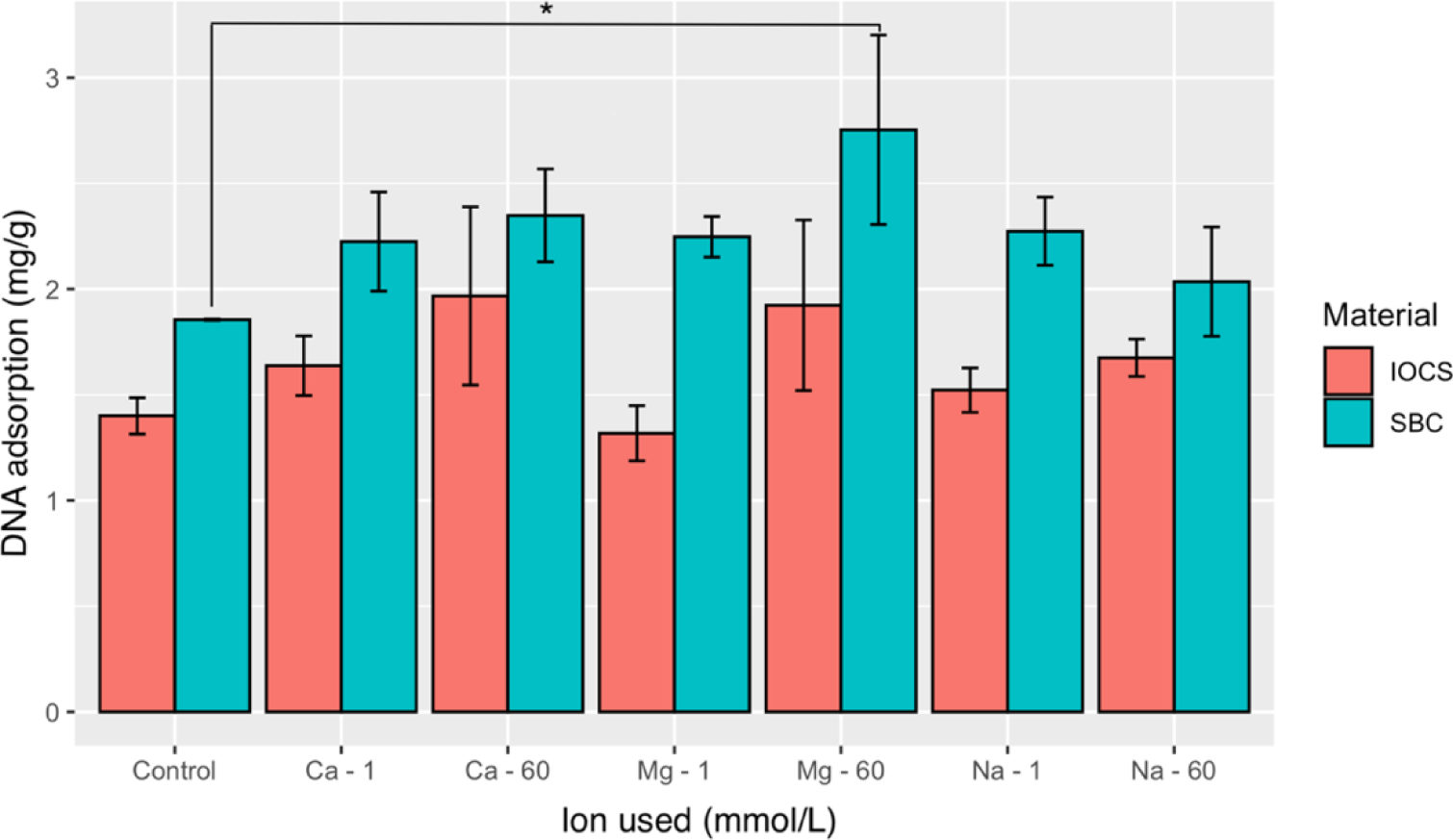
The influence of Ca^+2^, Mg^2+^ and Na^+^ on adsorption of DNA (100 mg L^-1^) on sewage sludge biochar (SBC) and iron oxide coated sands (IOCS). DNA adsorption stands for mg of DNA per g of adsorbent added. The concentrations of the ions were set at 1 and 60 mmol L^-1^. Significant difference is defined by (*) when p<0.05.

**Table 4.**
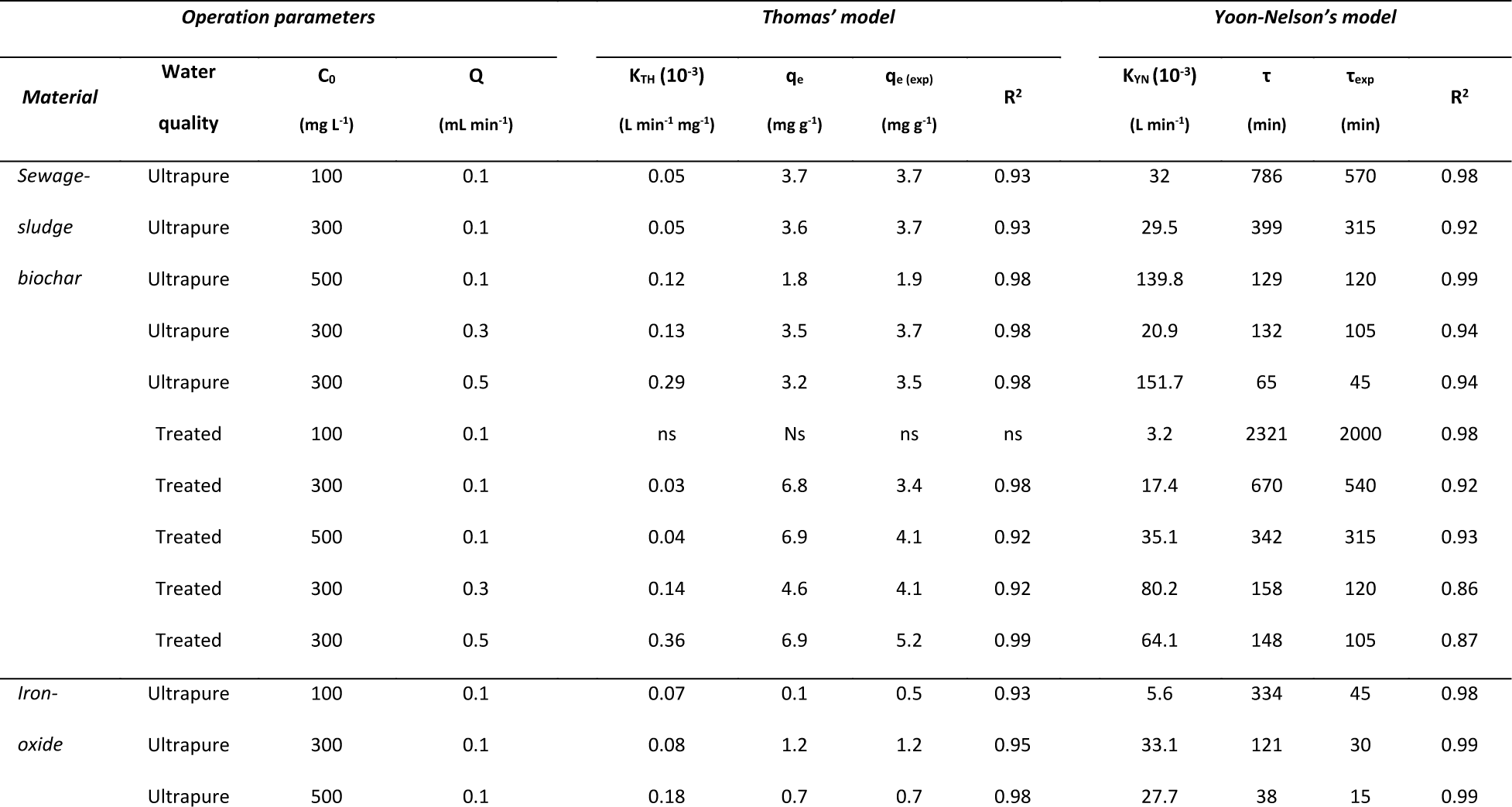
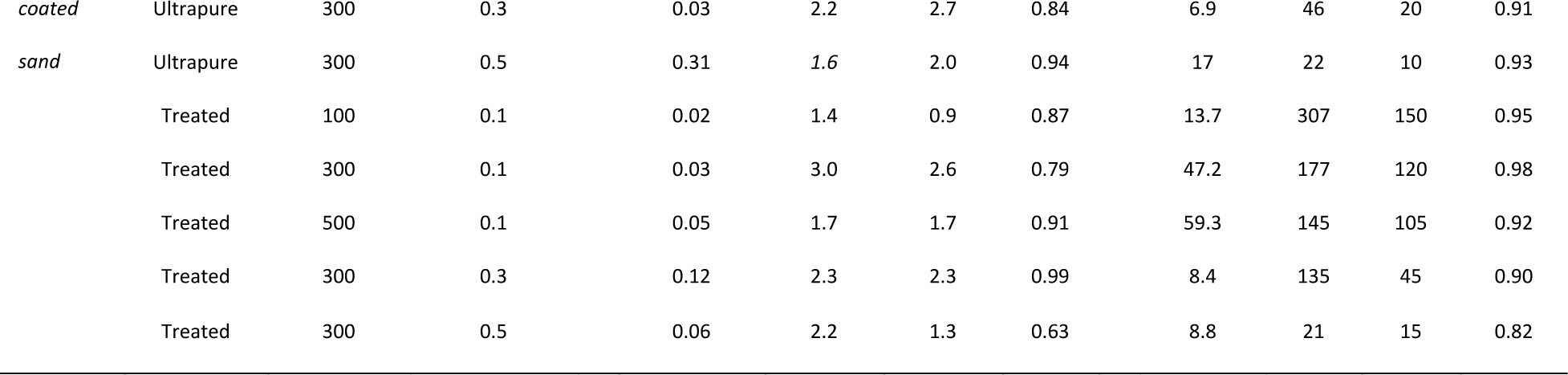
Parameters obtained from the Thomas and Yoon-Nelson models. **Ns:** non-saturated.

### 3.4 Sewage-sludge biochar is an efficient adsorbent for DNA removal

Two separate columns filled with sewage-sludge biochar or iron-oxide coated sand particles were evaluated as adsorbents for xenogenetic elements removal. Experiments were performed both with effluent and ultrapure water with salmon sperm DNA in order to assess the effect of treated wastewater on DNA adsorption.

The breakthrough curves from experiments performed with both columns and water qualities are depicted in **Figure 6.** Sewage-sludge biochar was more effective on DNA adsorption than iron oxide coated sand.

**Figure 6.**
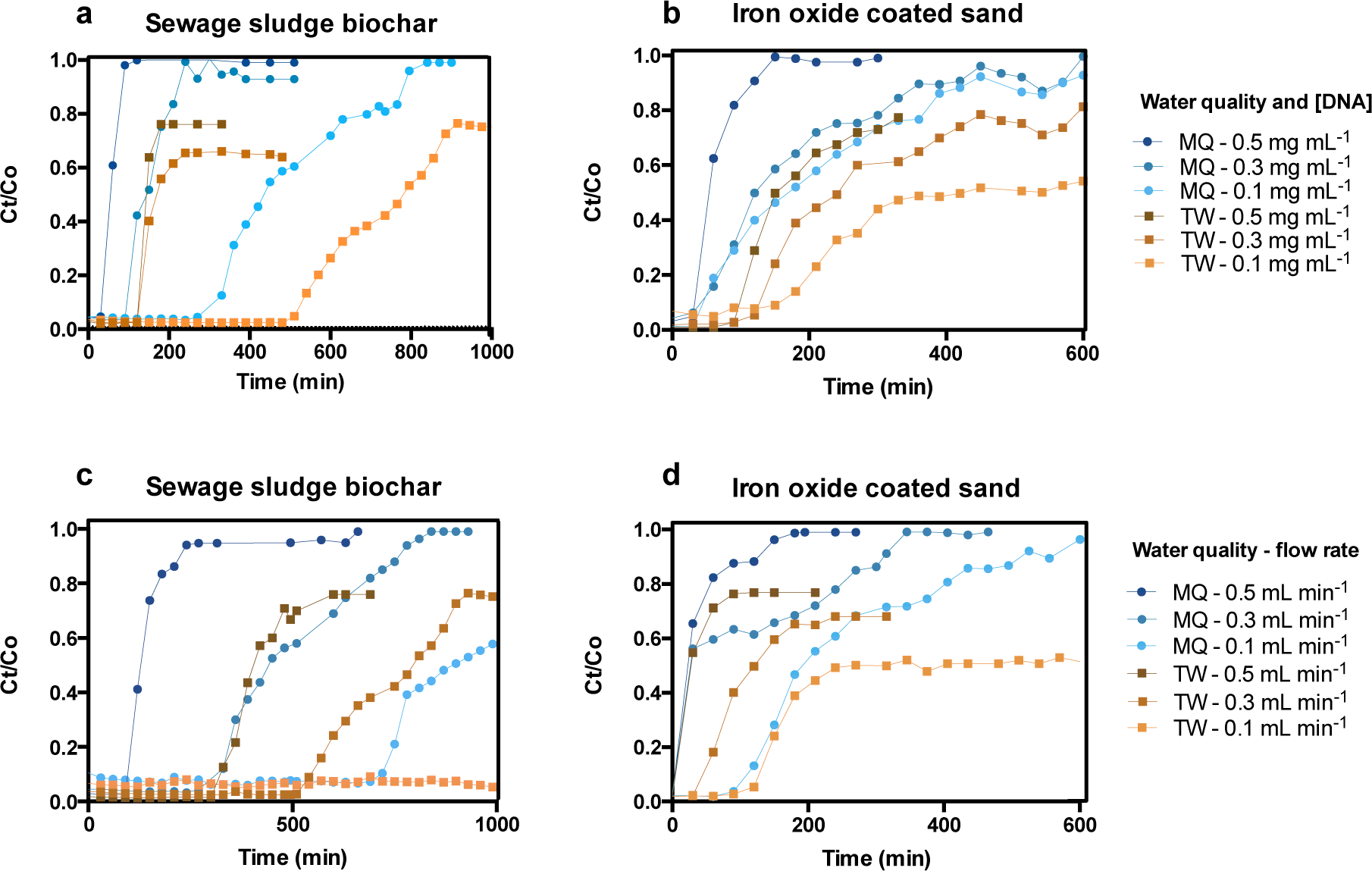
Comparison of experimental breakthrough curves for adsorption of salmon sperm DNA onto the different selected materials. Different initial concentrations ranging from 0.1 to 0.5 mg mL^-1^ and flow rates ranging from 0.1 to 0.5 mL min^-1^ were assessed on **(a)(c)** sewage sludge biochar and **(b)(d)** iron oxide coated sand, respectively. For assessing the effect of different initial DNA concentrations, flow rate was set at 0.1 mL min^-1^. For assessing the effect of different flow rates, the DNA concentration was set up to 0.3 mg mL^-1^. **MQ:** Ultrapure MiliQ water**. TW:** Treated wastewater.

The breakthrough point on sewage-sludge biochar was around 4.2 ± 1.8 times higher than iron oxide coated sand when different initial DNA concentrations were tested. The breakthrough point on sewage-sludge biochar was 4.6 ± 1.3 times higher than iron oxide coated sand when different flow rates were utilized.

When the same experiment was performed using ultrapure water, the effect on the breakthrough point was remarkable. The breakthrough point on sewage-sludge biochar was around 10.4 ± 1.9 times higher than iron oxide coated sand when different initial DNA concentrations (0.1, 0.3 and 0.5 mg mL^-1^) were tested and 4.3 ± 0.9 that of iron oxide coated sand when different flow rates (0.1, 0.3 and 0.5 mL min^-1^) were utilized. Full breakthrough point details can be found on **Table S5**.

**Table 5.**
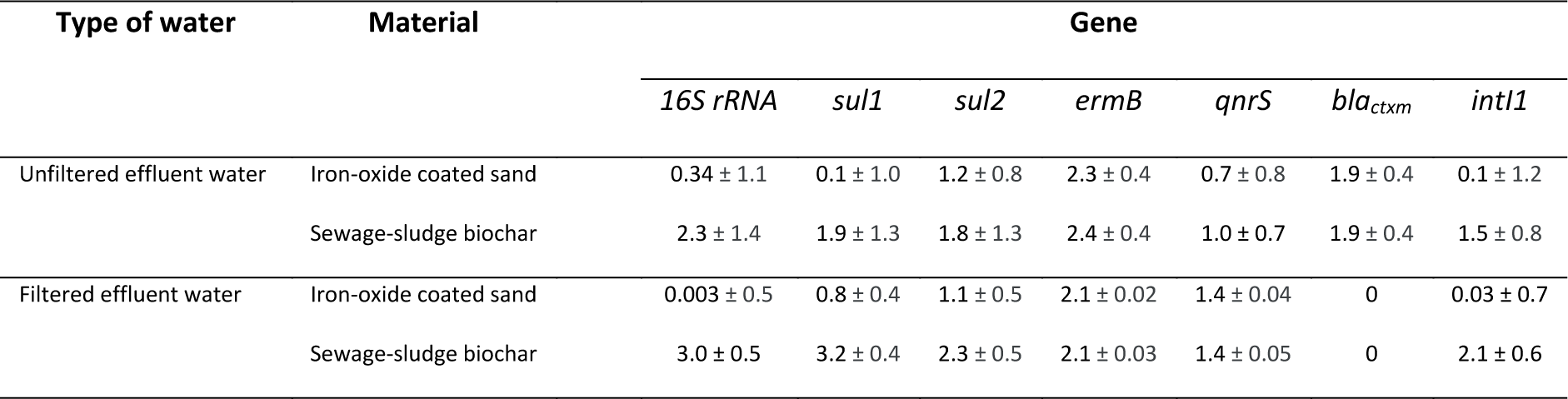
Quantitative PCR results assessing a 16S rRNA, a panel of antibiotic resistance gene and the integrase I class 1 as a mobile genetic element. Values represent the log10 gene copies differences between the inlet of the column (unfiltered or filtered effluent water) and the outlet of the adsorption columns tested (iron-oxide coated sand and sewage-sludge biochar).

| Type of water | Material | Gene |  |  |  |  |  |  |
| --- | --- | --- | --- | --- | --- | --- | --- | --- |
|  |  | 16S rRNA | *sul1* | *sul2* | *ermB* | *qnrS* | *bla<sub>ctxm</sub>* | *int11* |
| Unfiltered effluent water | Iron-oxide coated sand | 0.34 ± 1.1 | 0.1 ± 1.0 | 1.2 ± 0.8 | 2.3 ± 0.4 | 0.7 ± 0.8 | 1.9 ± 0.4 | 0.1 ± 1.2 |
|  | Sewage-sludge biochar | 2.3 ± 1.4 | 1.9 ± 1.3 | 1.8 ± 1.3 | 2.4 ± 0.4 | 1.0 ± 0.7 | 1.9 ± 0.4 | 1.5 ± 0.8 |
| Filtered effluent water | Iron-oxide coated sand | 0.003 ± 0.5 | 0.8 ± 0.4 | 1.1 ± 0.5 | 2.1 ± 0.02 | 1.4 ± 0.04 | 0 | 0.03 ± 0.7 |
|  | Sewage-sludge biochar | 3.0 ± 0.5 | 3.2 ± 0.4 | 2.3 ± 0.5 | 2.1 ± 0.03 | 1.4 ± 0.05 | 0 | 2.1 ± 0.6 |

Thomas’ and Yoon-Nelson’s mathematical models were used to evaluate the effect of process variables on the efficiency of adsorption for DNA removal in a fixed-bed column.

Breakthrough curves for the adsorption of DNA onto sewage-sludge biochar and iron-oxide coated sand on different water qualities were further analyzed using the Thomas model. The Thomas model constant “KTH” values increased while flow rates increased on both tested materials and water qualities. The adsorption capacity at equilibrium “qe” values calculated from the Thomas model were closer to the experimentally obtained results (qe (exp)), especially on ultrapure water. Thus, the Thomas model adequately describes the experimental breakthrough data. The suitability of the Thomas model to the experimental data indicates that external and internal diffusions are not the only rate-limiting steps in the DNA adsorption process.

The Yoon-Nelson model was applied to determine the extent of the adsorbents used for DNA removal. The model constants (KYN and τ) and correlation coefficient values are also presented in **table 4** for all experimental conditions. The interpolation of the model plot showed that the values of KYN increased when increasing inlet DNA concentrations. The breakthrough time ‘τ’ decreased for the increasing range of flow rate and initial DNA concentrations, as the column saturated faster due to less contact time and higher number of DNA molecules to be adsorbed. The small differences between experimental and predicted τ values indicated that Yoon-Nelson model gave an appropriate fit to the experimental column data on continuous DNA adsorption.

### 3.5 Humic acids interfere on DNA adsorption with effluent water matrices

For the ultrapure water experiments, both sewage-sludge biochar and iron-oxide coated sand were fully saturated by salmon sperm DNA (ratio Ct/Co≈1). This exhaustion point could not be achieved when working with treated wastewater **(Figure 6).** Such differences on different water qualities could be explained by DNA being adsorbed on some organic particles or components of treated wastewater that are not present on ultrapure water such as humic acids. If DNA binds to organic particles and do not precipitate, they will remain in solution and would not be detected in the eluent.

Humic acids are generally seen as important soil and natural water components that are formed during humification of organic matter by microorganisms. They are recognized as responsible for binding major parts of the available metal ions in water and soil (Kochany and Smith, 2001). A range of humic acid concentrations (0 to 100 mg L^-1^) were incubated with salmon sperm DNA and the adsorbents in order to assess the influence of humic acids on DNA adsorption.

Increasing humic acid concentrations decreased significantly the capacity of the materials tested in this study to adsorb DNA onto sewage-sludge biochar (Δ0.5 mg g^-1^, 90.5% decrease) and iron-oxide coated sand (Δ0.12 mg g^-1^, 17% decrease) (**Figure 7**). When only humic acids were supplied, DNA adsorbed onto them (0.42 mg g^-1^). Saeki *et al,* (2011) already showed a similar effect when DNA molecules were exposed to humic acids, suggesting that humic acids can adsorb and fix DNA. Humic acids can adsorb themselves onto biochar (Feng et al., 2008), binding better DNA and in consequence making biochar a better adsorbent. This would explain why the column outlet concentration with treated wastewater never reached the inlet DNA concentration when saturated.

**Figure 7.**
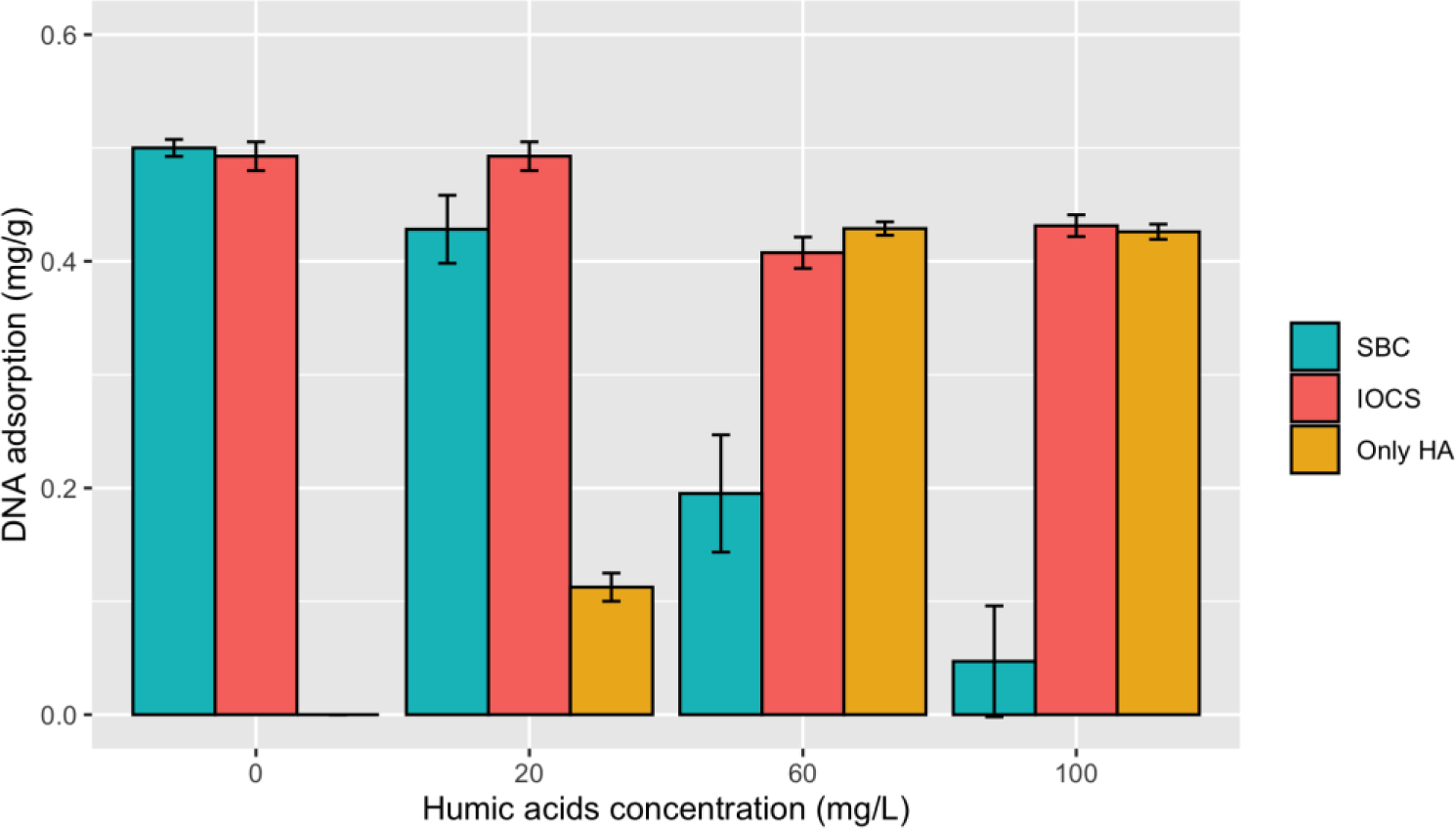
Influence of humic acids on DNA adsorption on sewage sludge biochar (SBC), iron oxide coated sands (IOCS) and only humic acids (HA). Salmon sperm DNA was used as a DNA template at a concentration of 20 µg mL^-1^. DNA adsorption stands for mg of DNA per g of adsorbent added.

### 3.6 More than 95% of antibiotic resistant genes and mobile genetic element present in the environmental DNA can be removed by sewage-sludge biochar

To assess the removal of ARGs and MGE, two columns with the adsorbents were used to treat 1000 mL of treated wastewater. To differentiate between ARGs and MGE removal in the environmental DNA or free-floating extracellular DNA, non-filtered and 0.2-µm filtered treated wastewater was used as mobile fraction.

The removal effect for a specific set of ARGs and MGE was assessed by qPCR. The results for the filtration experiments with the raw effluent are shown in **Figure 8a** and **Table 5**. The analysis of variance (ANOVA) on these scores yielded significant variation among the non-filtered effluent water and the eluent of the iron-oxide coated sand and sewage-sludge biochar columns. The post-hoc Tukey test showed that all the tested gene copies in the sewage-sludge biochar eluent differed significantly at p<.0005 when compared with their concentration on the non-filtered effluent wastewater inlet. For the iron coated sand, only *sul2, ermB,* and *blaCTXM* differed significantly at p<0.0005 and *qnrS* at p<0.05 from the inlet concentrations. Individual qPCR gene values per sample are listed in **Table S6**.

**Figure 8.**
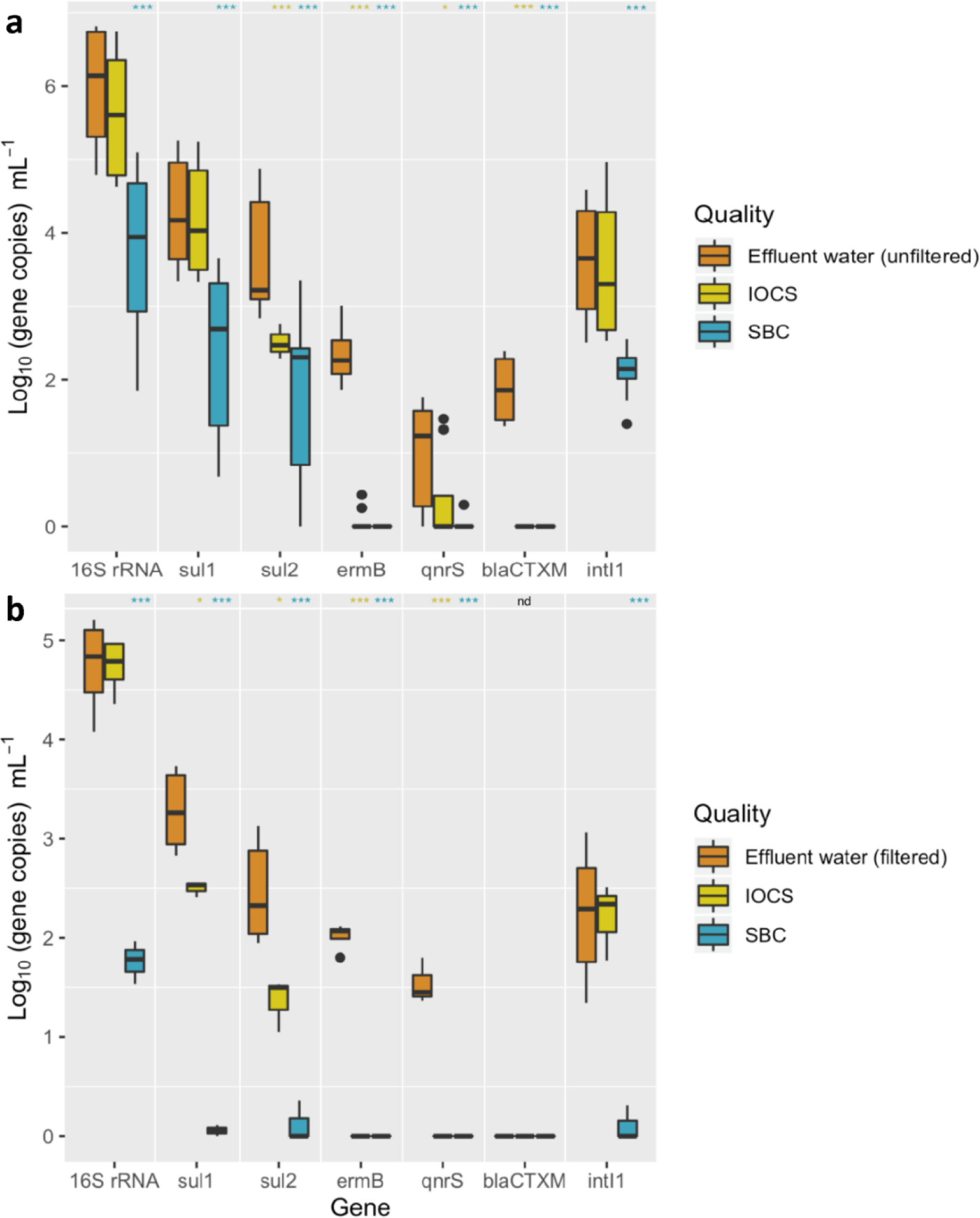
Quantitative PCR results assessing the ARGs and MGE differences between treated effluent wastewater and the by-product column outlet water streams after running through the iron-oxide-coated sand (IOCS) adsorption column or the sewage-sludge biochar (SBC) adsorption column. **(a)** Comparison is done with unfiltered effluent water samples assessing the environmental DNA removal (including antibiotic resistance bacteria). **(b)** Comparison is done with filtered effluent water samples assessing the free-floating extracellular DNA removal (only including antibiotic resistance genes and mobile genetic elements. No bacteria presence). Values are shown on Log10 gene copies mL^-1^ from a panel of 16S rRNA, different antibiotic resistance genes (*sul1, sul2, ermB, qnrS and blaCTXM*) and a mobile genetic element (*intI1*). p<0.05(*), p<0.005 (**), p<0.0005 (***). **Note:** “nd” stands for “non-detected”. Black dots represent data outliers.

The results from filtered effluent wastewater and the sewage-sludge biochar and iron-oxide coated sand column eluents are shown in **Figure 8b** and **table 5**. An ANOVA on these scores yielded significant variation among the filtered effluent water and the eluent of the iron-oxide coated sand and sewage-sludge biochar columns. No significant difference between samples were observed when *blaCTXM* was assessed, basically because it was not detected (n.d.) in any fraction. The post-hoc Tukey test showed that all the tested gene copies in the sewage-sludge biochar eluent differed significantly from the filtered treated wastewater at p<.0005. Only

*sul1* and *sul2* (p<0.05) and *ermB* and *qnrS* (p<0.0005) differed significantly from the iron-oxide coated sand eluent when compared with the column inlet concentrations. Individual qPCR gene values per sample are listed in **Table S7**.

Overall, 97% and 66% of the genes present in the unfiltered effluent water decreased in these experiments using the sewage-sludge biochar columns and iron-oxide coated sand, respectively. When the free-floating extracellular DNA was assessed, a removal of 84.8% and 54% for sewage-sludge biochar columns and iron-oxide coated sand columns was observed, respectively. Individual removal values are listed in **Table S8**.

In this study, other materials such as granulate activated carbon, particulate activated carbon and mineral wool were tested on top of the selected materials (**Figure S4**). Only the particulate activated carbon, used as positive control, the sewage-sludge biochar and the iron-oxide coated sands displayed positive results on DNA adsorption. This is the first time to our knowledge that wastewater treatment by-products have been shown to be effective adsorbents for antibiotic resistance genes and mobile genetic elements from the free-floating extracellular DNA fraction removal from treated effluent water.

### 3.7 Outlook

Sewage-sludge biochar can be a good by-product option for ARGs and MGEs removal by adsorption. Sewage-sludge has been gradually increasing due to the rising world population. It is estimated that at the European level, sludge production from WWTPs would reach 13 million tons by 2020 (Capodaglio and Callegari, 2018). The treatment and discarding of sewage sludge (landfilling, agriculture and incinerations) is an expensive and ecological burden due to urban expansion. This causes an increase in sludge production as new WWTPs are required to be built. The European directive 86/278/EEC on agricultural use of sewage sludge set stringent regulations on the use of sewage sludge as landfilling due to the presence of high concentrations of heavy metals and pathogens. Incineration is carried out in most of the EU-15 countries, with the Netherlands one of the top users (Agrafioti et al., 2013; Capodaglio and Callegari, 2018).

One of the options for getting rid of the sewage sludge is large scale incineration, which requires high investment and operating costs as extensive cleaning, as well as gas purification for safe emission into the environment. An alternative to this method would be pyrolysis which can reduce the sludge volume, completely remove pathogens and convert the organic matter into biofuel, bio-oil or biochar for further applications (Inguanzo et al., 2002). Pyrolysis is done under less or no oxygen conditions which reduces the amount of flue gases to be cleaned and acidic gases generation (Hwang et al., 2007). Biochar produced out of dewatered sewage-sludge has been shown to be useful in removing pollutants from wastewater (Fathi Dokht et al., 2017). The ability to remove nucleic acids is an additional advantage of this by-product material on top of removing agrichemicals, pharmaceuticals and personal care products as well as endocrine disrupting compounds (Thompson et al., 2016).

Methods for removing ARBs and ARGs from effluent water have so far primarily focused on killing bacteria rather than removing the DNA. Conventional disinfectants such as chlorine or ozone, UV treatments or advanced oxidation processes (fenton oxidation or photocatalytical oxidation, among others) have been used (Pazda et al., 2019). We show here an alternative to non-mutagenic treatments, simpler and cost-effective.

A free-floating eDNA concentration of 5.6 ± 2.9 µg L^-1^ has been measured from effluent wastewater (Calderón-Franco et al., 2020b). The WWTP sampled in this study has an approximate flow of 200’000 m^3^ day^-1^, corresponding to an approximate free-floating extracellular DNA discharge of 1 kg day^-1^. If we take from this study that sewage-sludge biochar is efficient to remove around 85% of the free-floating genes, it would mean that we could avoid discharging 310 kg year^-1^ of it from only one, but also the largest in the country, Dutch WWTP.

## 4 Conclusion

We pointed out the nucleic acids removal capacity of water sanitation by-products, highlighting the importance of giving them a second productive life. This was remarkable for sewage-sludge biochar as it can be combined with organic micropollutants removal. We showed with our laboratory physical models how to efficiently remove ARGs and MGEs before water leaves the WWTP. The 97% of the evaluated genes present in the environmental DNA (composed of both the intracellular and extracellular DNAs) were adsorbed and removed from raw unfiltered effluent wastewater by sewage-sludge biochar; 66% by iron-oxide coated sand. The 85% of ARGs and MGE present on free-floating eDNA were removed by sewage-sludge biochar; 54% by iron-oxide coated sand. The use of sewage-sludge biochar in combination with humic acids found in treated effluents can impede the release of around 310 kg day^-1^ of free-floating eDNA into the environment. By aligning to the UN SDGs for clean water and sanitation, responsible consumption and production, and climate action, this study provides a definite solution for the remediation of xenogenetic elements from wastewater. Both end-of-pipe and decentralized technologies can be developed to prevent their discharge in the aquatic environment and to minimize their emission at the source, respectively.

## Authors’ contributions

DCF and AS designed the study with DGW, and additional input from MvL on adsorbent selection. DCF and AS performed the experimental investigations. DCF, AS, and DGW critically assessed the experiments, data, and scientific findings with periodical feedbacks from GJM. DCF and AS prepared the outline of the manuscript. DCF wrote the manuscript and crafted all figures with direct contribution, edits, and critical feedback by all authors.

## Acknowledgements

We are grateful to Suellen Espindola from the TU Delft Department of Chemistry for taking the SEM micrographies and to Dr. Iulian Dugulan from the TU Delft Reactor Institute for characterizing our iron-oxide coated sands by Mössbauer spectroscopy. We are also thankful to John van den Berg from the TU Delft Department of Material Sciences for his guidance through the nitrogen sorption analysis pipeline. This work is part of the research project “Transmission of Antimicrobial Resistance Genes and Engineered DNA from Transgenic Biosystems in Nature” (Targetbio) funded by the programme Biotechnology & Safety (grant no. 15812) of the Applied and Engineering Sciences Division of the Dutch Research Council (NWO). This manuscript will be made available as a pre-print on bioRxiv.

## Supplementary information

### qPCR mix solution and reaction conditions

All ARGs and *intI1* qPCR reactions were conducted in 20 µL, including IQ^TM^ SYBR green supermix BioRad 1x. Forward and reverse primers, and oligonucleotide probes (when applicable) are summarized in **Table S1 and S2.** A total of 2 µL of DNA template was added to each reaction, and the reaction volume was completed to 20 µL with DNase/RNase free Water (Sigma Aldrich, UK). All reactions (were performed in a qTOWER3 Real-time PCR machine (Westburg, DE) according to the following PCR cycles: 95°C for 5 min followed by 40 cycles at 95°C for 15 s and 60°C for 30 s. The annealing temperature was the same for all the different reactions except for the s*ul2* and *sul1* genes. In those cases, the annealing temperatures were 61°C and 65°C, respectively.

In order to check the specificity of the reaction, a melting curve was performed from 65 to 95°C at a temperature gradient of +0.5°C (5 s)^-1^. Synthetic DNA fragments (IDT, USA) containing each of the target genes were used as a positive control to create the standard curves. Serial dilutions of gene fragments were performed in sheared salmon sperm DNA 5 µg mL^-1^ (m/v) (Thermofisher, LT) diluted in Tris-EDTA (TE) buffer at pH 8.0. Every sample was analyzed in technical triplicates. Standard curves were included in each PCR plate with at least 6 serial dilutions points and in technical duplicate. An average standard curve based on a standard curve from every run was created for every gene set. Gene concentration values were then calculated from the aforementioned curve.

**Table S1.**
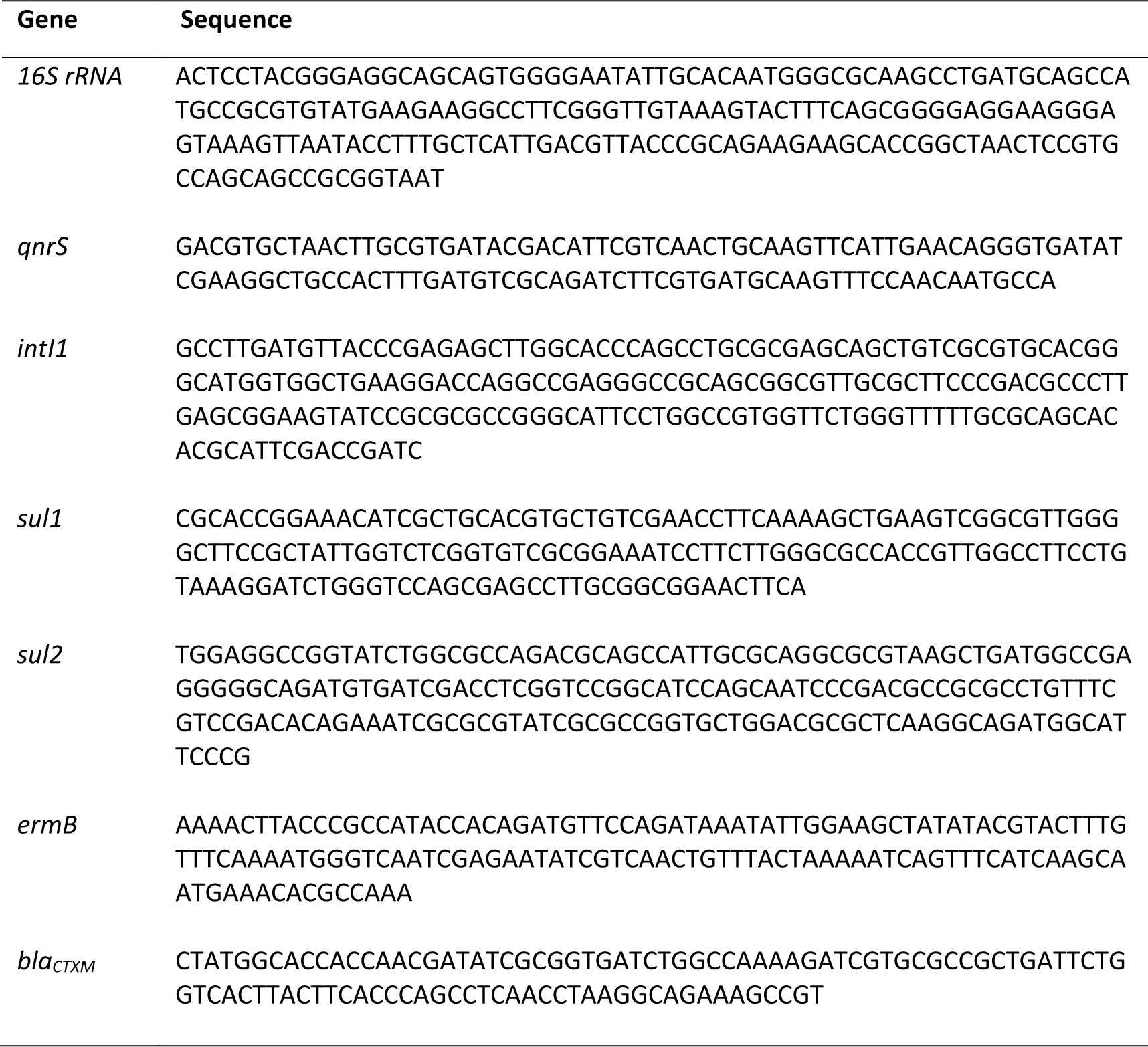
16S rRNA, ARGs and MGE synthetic DNA fragments used from ResFinder for generating standard curves used for qPCR analysis.

**Table S2.**
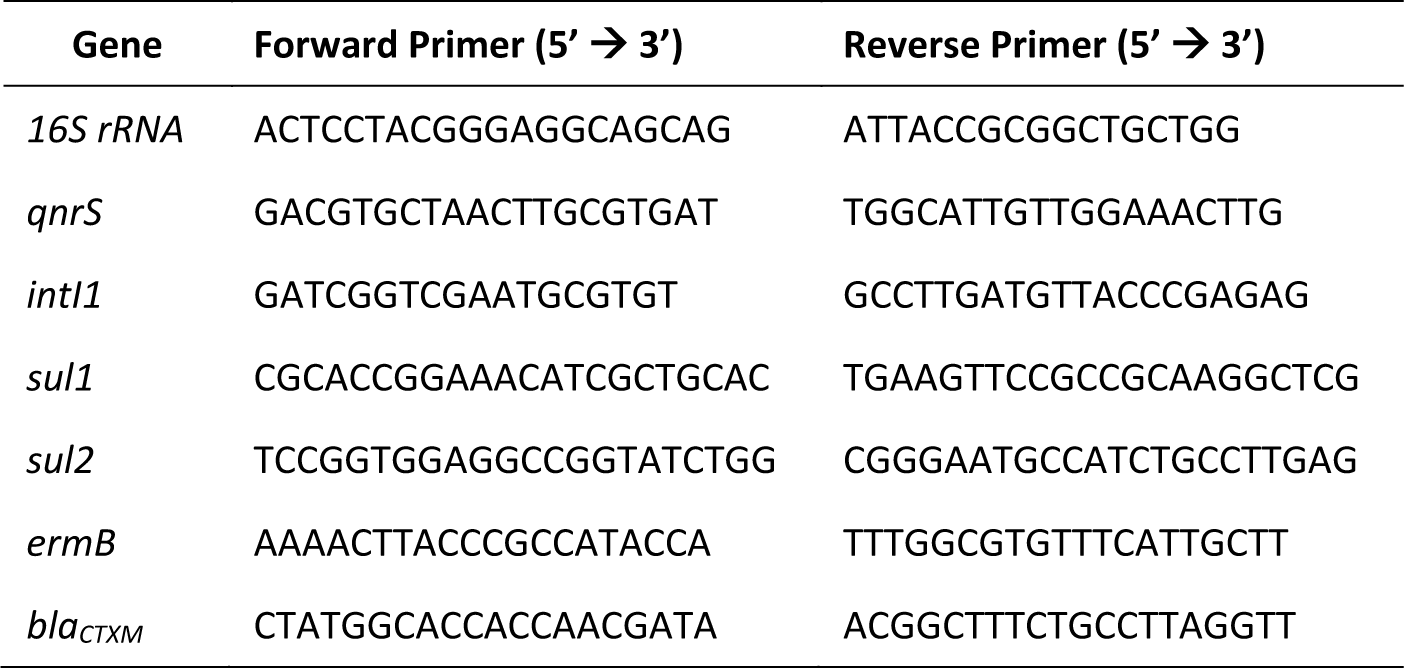
Primers used in this study.

### Mössbauer spectra and fitted parameters for characterization of iron-oxide coated sand

**Figure S1.**
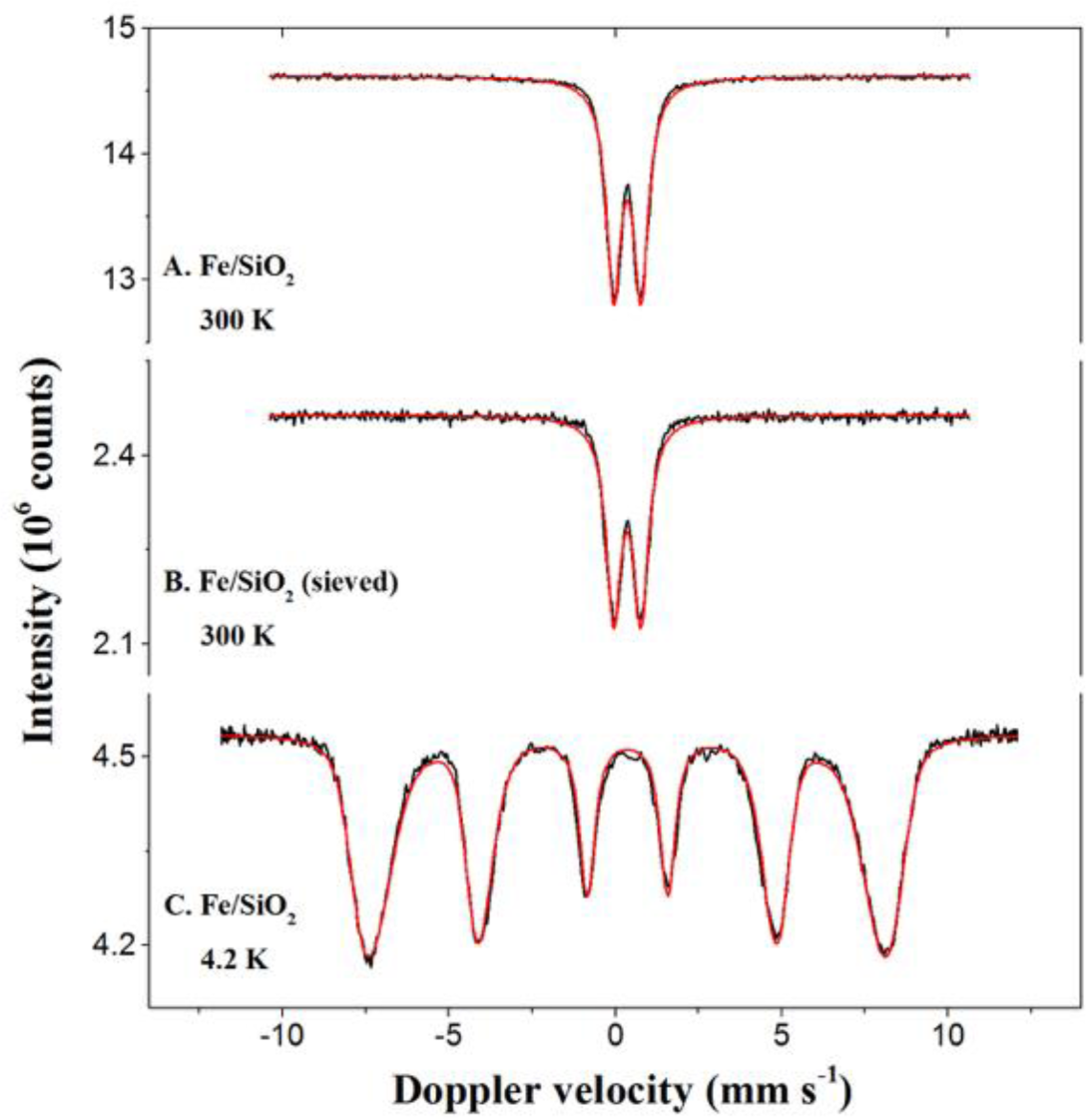
Mössbauer spectra obtained at 300 and 4.2 K with the Fe/SiO2samples.

**Table S3.** The Mössbauer fitted parameters of the Fe/SiO2samples.

| Sample | T<br>(K) | IS<br>(mm·s<sup>-1</sup>) | QS<br>(mm·s<sup>-1</sup>) | Hyperfine<br>field (T) | Γ<br>(mm·s<sup>-1</sup>) | Phase | Spectral<br>contribution (%) |
| --- | --- | --- | --- | --- | --- | --- | --- |
| Fe/SiO<sub>2</sub> | 300 | 0.36 | 0.82 | -<br>- | 0.53 | Fe<sup>3+</sup> | 100 |
| Fe/SiO<sub>2</sub><br>(sieved) | 300 | 0.36 | 0.82 | -<br>- | 0.53 | Fe<sup>3+</sup> | 100 |
| Fe/SiO<sub>2</sub> | 4.2 | 0.35 | -0.02 | 47.4* | 0.48 | Fe<sup>3+</sup><br>(Ferrihydrite) | 100 |

Experimental uncertainties: Isomer shift: I.S. $\pm 0.01$ mm s<sup>-1</sup>; Quadrupole splitting: Q.S. $\pm 0.01$ mm s<sup>-1</sup>; Line width: $\Gamma \pm 0.01$ mm s<sup>-1</sup>; Hyperfine field: $\pm 0.1$ T; Spectral contribution: $\pm 3\%$ ; \*Average magnetic field.

### By-product materials macroscopic visualization and setup used for the experiments

**Figure S2.**
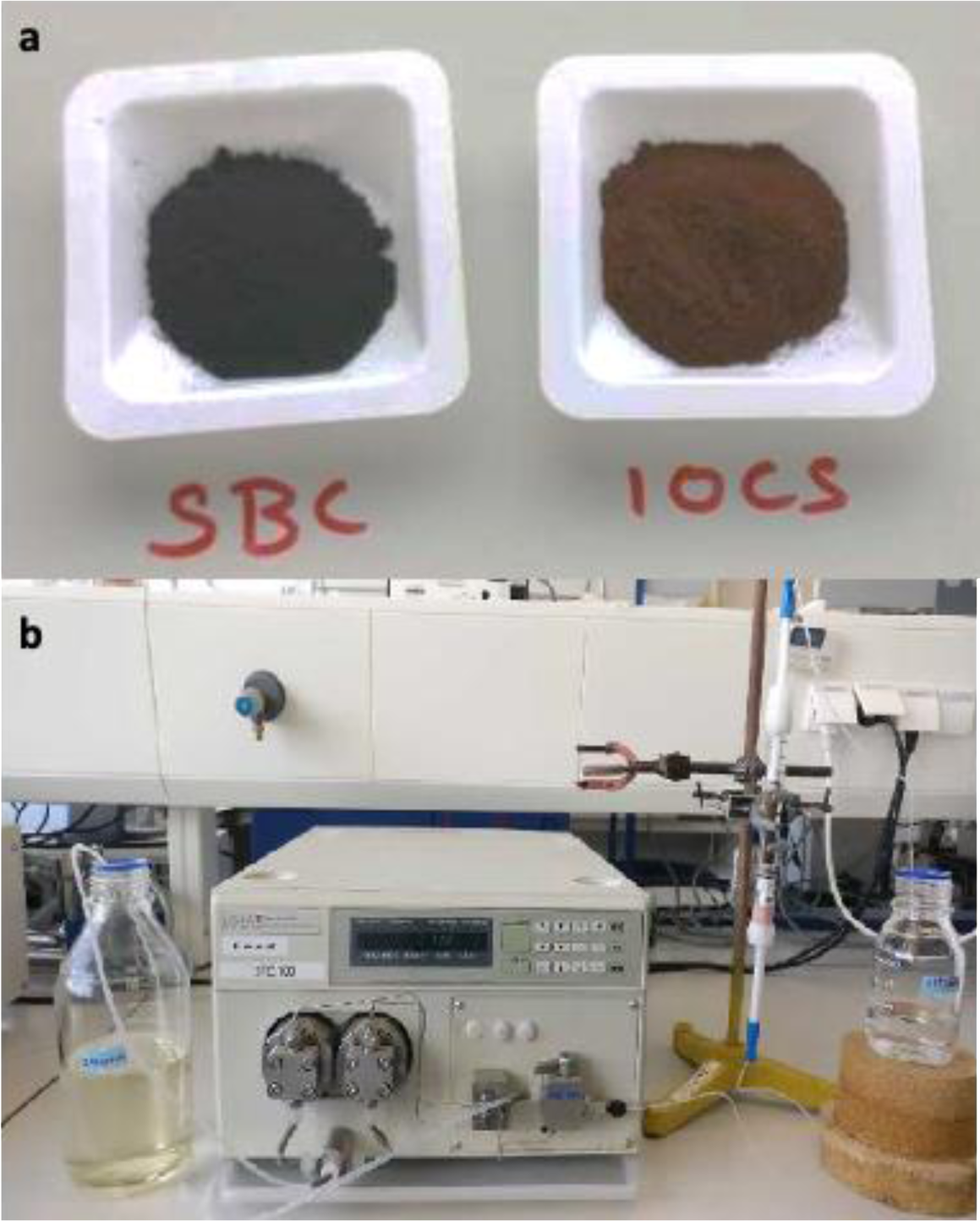
By-product materials macroscopic visualization (a) and setup used for the column-based studies (b).

### Equilibrium adsorption time from both by-product materials

**Figure S3.**
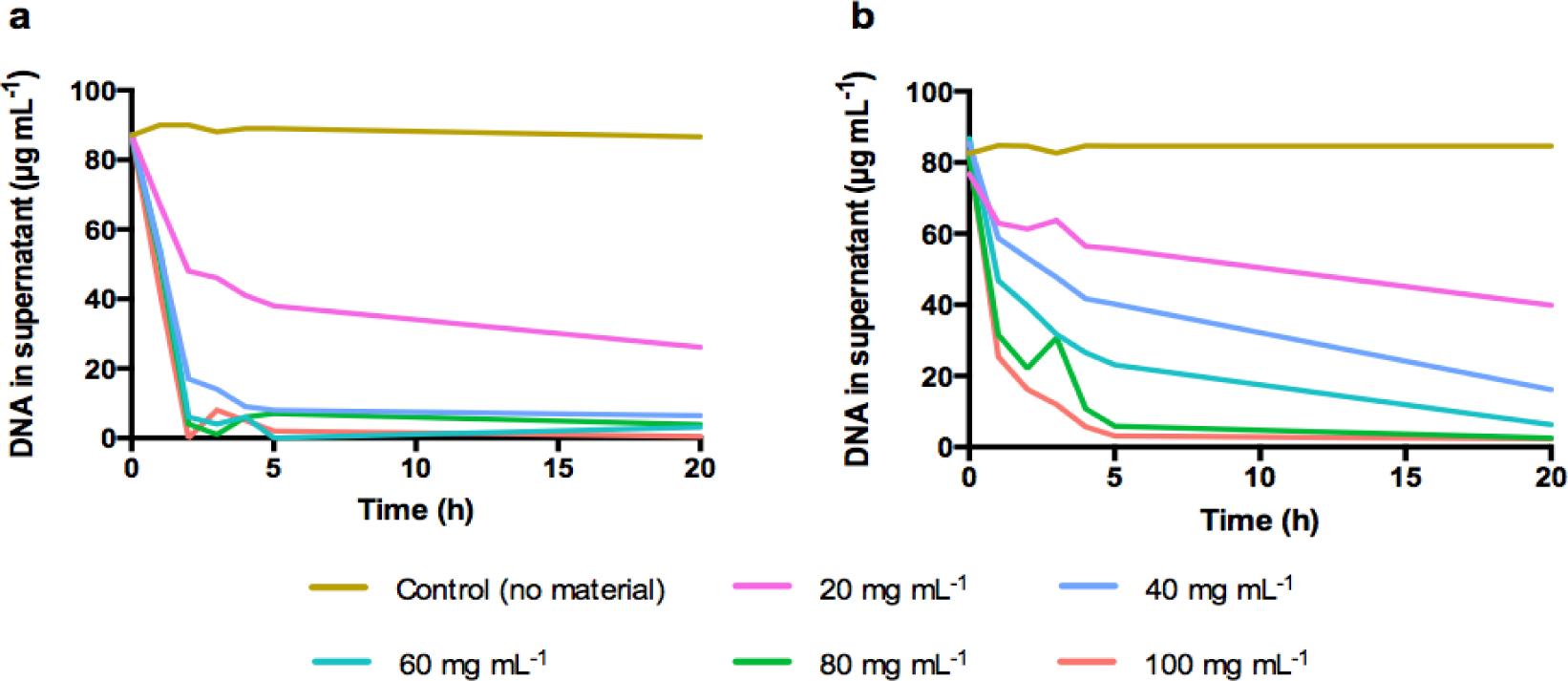
Equilibrium time on treated wastewater for **(a)** sewage sludge biochar and **(b)** iron oxide coated sands. Initial DNA concentration of 100 µg mL^-1^ with the adsorbent ranging from 0 to 100 mg mL^-1^.

**Table S4.** Statistical significance from the comparisons done between the control condition (ultrapure water) and different ions concentrations (1 – 60 mM)

| Material | Comparison | diff | lwr | upr | p.adj |
| --- | --- | --- | --- | --- | --- |
| SBC | Control - Ca - 1 mM | -0.37 | -1.04 | 0.30 | 0.52 |
| SBC | Control - Ca - 60 mM | -0.49 | -1.16 | 0.18 | 0.23 |
| SBC | Control - Mg - 1 mM | 0.39 | -0.28 | 1.06 | 0.46 |
| SBC | Control - Mg - 60 mM | 0.90 | 0.23 | 1.57 | 0.01 |
| SBC | Control - Na - 1 mM | 0.42 | -0.25 | 1.09 | 0.39 |
| SBC | Control - Na - 60 mM | 0.18 | -0.49 | 0.85 | 0.96 |
| IOCS | Control - Ca - 1 mM | -0.23 | -0.91 | 0.43 | 0.88 |
| IOCS | Control - Ca - 60 mM | -0.57 | -1.24 | 0.10 | 0.12 |
| IOCS | Control - Mg - 1 mM | -0.08 | -0.75 | 0.59 | 0.99 |
| IOCS | Control - Mg - 60 mM | 0.52 | -0.15 | 1.19 | 0.18 |
| IOCS | Control - Na - 1 mM | 0.12 | -0.55 | 0.79 | 0.99 |
| IOCS | Control - Na - 60 mM | 0.27 | -0.39 | 0.94 | 0.79 |

**Table S5.** Adsorbents breakthrough points depending on the flow rate and DNA concentration

| Water quality | Adsorbent | Flow rate<br>(mL min<sup>-1</sup>) | DNA<br>concentration<br>(mg L<sup>-1</sup>) | Breakthrough<br>point<br>(min) |
| --- | --- | --- | --- | --- |
| Treated wastewater | Biochar | 0.1 | 0.3 | 510 |
|  |  | 0.3 |  | 120 |
|  |  | 0.5 |  | 105 |
|  | IOCS | 0.1 | 0.3 | 120 |
|  |  | 0.3 |  | 45 |
|  |  | 0.5 |  | 15 |
|  | Biochar | 0.1 | 0.1 | >2000 |
|  |  |  | 0.3 | 540 |
|  |  |  | 0.5 | 315 |
|  | IOCS | 0.1 | 0.1 | 150 |
|  |  |  | 0.3 | 120 |
|  |  |  | 0.5 | 105 |
| Ultrapure water | Biochar | 0.1 | 0.3 | 315 |
|  |  | 0.3 |  | 105 |
|  |  | 0.5 |  | 45 |
|  | IOCS | 0.1 | 0.3 | 105 |
|  |  | 0.3 |  | 20 |
|  |  | 0.5 |  | 10 |
|  | Biochar | 0.1 | 0.1 | 570 |
|  |  |  | 0.3 | 315 |
|  |  |  | 0.5 | 120 |
|  | IOCS | 0.1 | 0.1 | 45 |
|  |  |  | 0.3 | 30 |
|  |  |  | 0.5 | 15 |

**Table S6.**
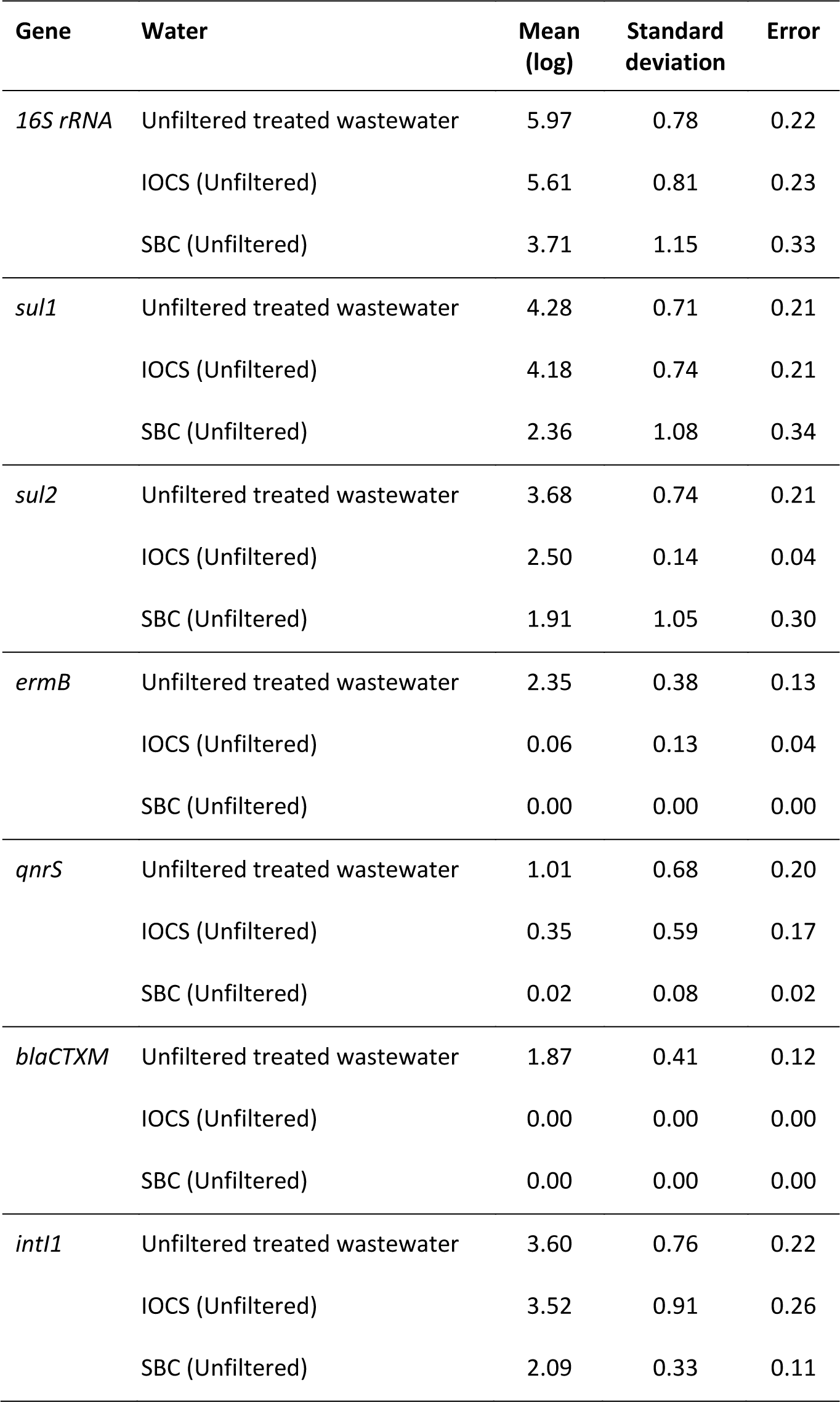
qPCR results per gene from unfiltered treated wastewater and eluents after IOCS and SBC columns

**Table S7.** qPCR results per gene from filtered treated wastewater and eluents after IOCS and SBC columns

| Gene | Sample | Mean (Log) | Standard deviation | Error |
| --- | --- | --- | --- | --- |
| *16S rRNA* | Filtered treated wastewater | 4.74 | 0.44 | 0.22 |
|  | IOCS (Filt.) | 4.74 | 0.23 | 0.10 |
|  | SBC (Filt.) | 1.76 | 0.18 | 0.10 |
| *sul1* | Filtered treated wastewater | 3.28 | 0.37 | 0.15 |
|  | IOCS (Filt.) | 2.50 | 0.06 | 0.04 |
|  | SBC (Filt.) | 0.05 | 0.05 | 0.04 |
| *sul2* | Filtered treated wastewater | 2.46 | 0.47 | 0.19 |
|  | IOCS (Filt.) | 1.36 | 0.22 | 0.13 |
|  | SBC (Filt.) | 0.12 | 0.17 | 0.10 |
| *ermB* | Filtered treated wastewater | 2.08 | 0.03 | 0.01 |
|  | IOCS (Filt.) | 0.00 | 0.00 | 0.00 |
|  | SBC (Filt.) | 0.00 | 0.00 | 0.00 |
| *qnrS* | Filtered treated wastewater | 1.41 | 0.04 | 0.03 |
|  | IOCS (Filt.) | 0.00 | 0.00 | 0.00 |
|  | SBC (Filt.) | 0.00 | 0.00 | 0.00 |
| *blaCTXM* | Filtered treated wastewater | 0.00 | 0.00 | 0.00 |
|  | IOCS (Filt.) | 0.00 | 0.00 | 0.00 |
|  | SBC (Filt.) | 0.00 | 0.00 | 0.00 |
| *int11* | Filtered treated wastewater | 2.23 | 0.61 | 0.25 |
|  | IOCS (Filt.) | 2.21 | 0.32 | 0.18 |
|  | SBC (Filt.) | 0.10 | 0.15 | 0.08 |

**Table S8.**
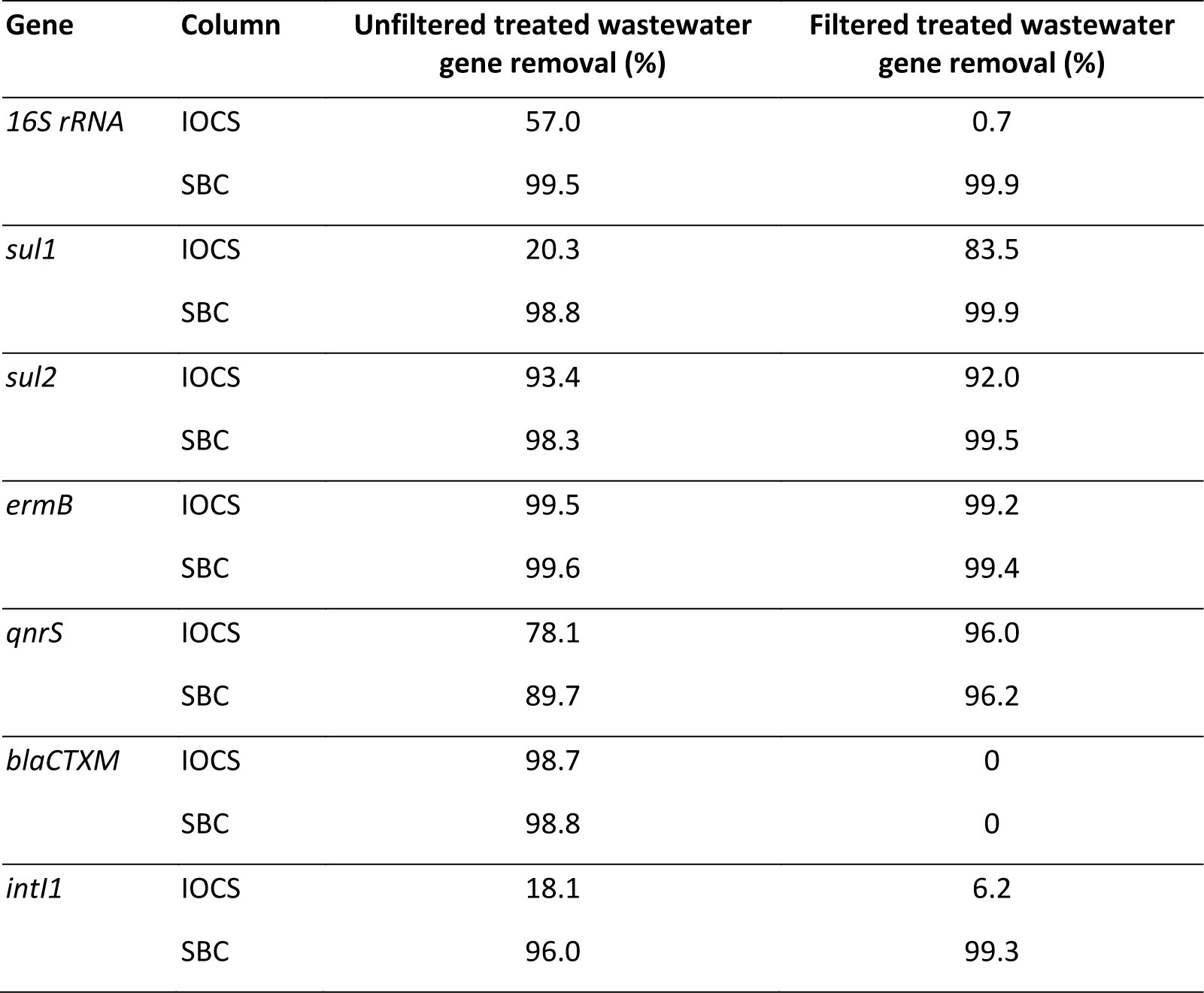
Gene removal (%) after filtered and unfiltered treated wastewater was run through IOCS and SBC columns.

### Preliminary experiments for selecting materials for DNA adsorption

**Figure S4.**
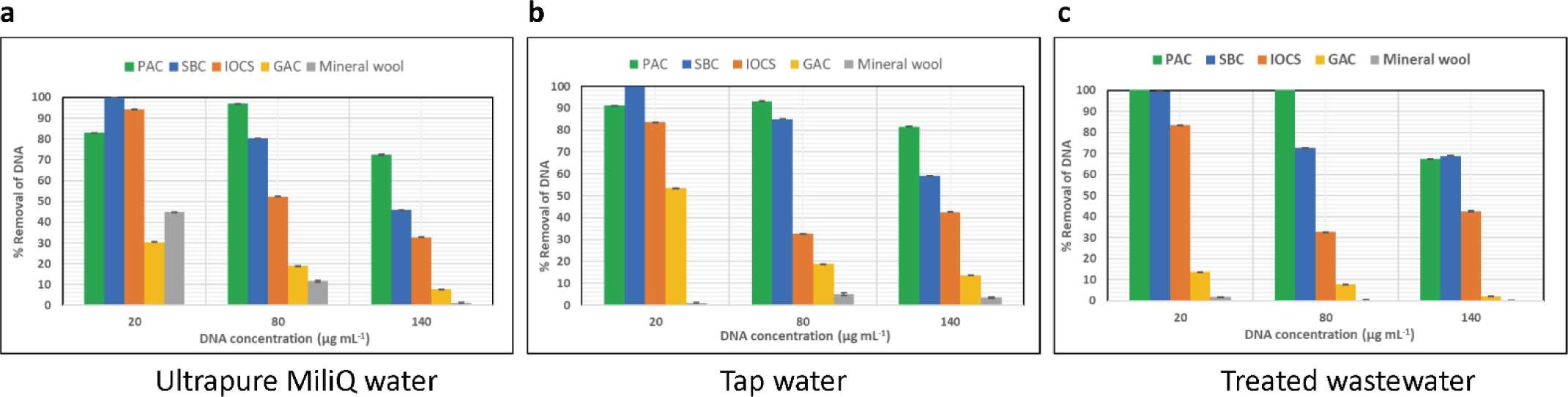
DNA removal in percentage with powdered activated carbon (PAC), sewage-based biochar (SBC), iron oxide coated sands (IOCS), granular activated carbon (GAC) and mineral wool in (a) ultrapure water (b) tap water (c) effluent wastewater.

### F-statistics and ANOVA

When compared non-filtered effluent water with IOCS and SBC eluents, F = 18.91, 15.72, 16.19, 347.1, 10.24, 230.2 and 12.06, p<0.0001 were observed for the panel of *16S rRNA, sul1, sul2, ermB, qnrS*, *blaCTXM* and *intI1* genes, respectively.

When comparing filtered effluent water with IOCS and SBC eluents, F = 78.0, 78.03, 31.9, 547, 76.2 and 17.6, p<0.0001 were observed for the panel of *16S rRNA, sul1, sul2, ermB, qnrS* and *intI1* genes, respectively.

